# Transitional lncRNA Signatures Reveal Distinct Stages of Cancer Progression and Metastasis

**DOI:** 10.1101/2025.06.10.658949

**Authors:** Stav Zok, Daniel Behar, Michal Linial

## Abstract

Long non-coding RNAs (lncRNAs) are emerging as key regulators in cancer, influencing gene expression, chromatin remodeling, and signaling. Evidence from The Cancer Genome Atlas (TCGA) and other datasets supports their role in tumor progression. Although the human genome harbors thousands lncRNA genes, only a small subset has been validated in cancer. In this study, we used the LncBook catalog (∼95,000 lncRNAs) to identify ∼12,500 lncRNAs with expression evidence across major TCGA cancer types. These were stratified by clinical annotations, including cancer stage (I–IV) and metastatic state (M0/M1). Using significant differential expression (z-score >|3|) for consecutive transitions, we identified a set of influential transitional lncRNAs (Tr-lncRNAs) that signify cancer transitions. Analyzing seven transitions revealed that over 70% of Tr-lncRNAs were cancer-type specific, while only 2–4% were shared across 17 major cancers. Each cancer type had 30–80 Tr-lncRNAs, with more than half uniquely expressed in one type. Most Tr-lncRNAs were previously uncharacterized. A pan-cancer analysis revealed 14 shared Tr-lncRNAs, including known ones such as XIST and H19. Our findings highlight distinct lncRNA expression patterns during cancer progression and provide new insights into cis-regulatory antisense mechanisms. We discuss the potential of Tr-lncRNAs as diagnostic biomarkers and therapeutic targets in cancer.

## Introduction

LncRNAs are defined as transcripts longer than 200 nucleotides without obvious protein-coding potential. Most lncRNA transcripts are lowly expressed and not evolutionarily conserved. It is estimated that approximately 1,000 human lncRNAs show signs of functionality and evolutionary conservation (Gao et al., 2020). A debate on the functional role of lncRNAs has yet to be resolved. The view that dominates lncRNAs argue that many of them are a byproduct of the transcriptional machinery (i.e., transcriptional “noise”) (Mattick et al., 2023). Another view claims that lncRNAs play a fundamental role in defining cellular states, development, and cell dynamics, particularly in affecting chromatin organization (Kopp and Mendell, 2018). Despite growing interest, distinguishing truly functional lncRNAs from noisy transcription remains a challenge (Perry and Ulitsky, 2016). LncRNAs have gained substantial attention for their roles in epigenetic regulation, chromatin dynamics, and post-transcriptional gene expression (Cao, 2014). An emerging notion is that the function of lncRNAs depends on their subcellular localization (in nuclei, exosomes) (Bridges et al., 2021), while regulating gene activity and cellular processes is dictated by their genomic position, capacity for miRNA sponging, and competition for cellular resources (e.g., transcription factors, RNA-binding proteins) (Yang et al., 2022). While the field of noncoding RNAs (ncRNAs) has flourished over the last four decades, biological understanding has mostly been associated with small ncRNAs (e.g., miRNAs, tRNAs, snoRNAs). Important examples of lncRNAs that play a fundamental role in development and cell fate include XIST, which drives X-chromosome inactivation in females, and H19, which is a maternally imprinted gene implicated in promoting cell growth, invasion, migration, epithelial-mesenchymal transition (EMT), metastasis, and apoptosis (Yang et al., 2021). Mechanistically, many lncRNAs may act as molecular sponges by sequestering certain miRNAs (Hashemi et al., 2022; Raveh et al., 2015). However, the low stoichiometry regarding their targets raises questions about their importance. Notably, the cumulative expression levels of a cell’s lncRNAs account for only a few percent of its transcriptome (Fatica and Bozzoni, 2014).

In conjunction with the maturation of deep sequencing techniques, the number of lncRNAs has rapidly grown, and the spectrum of methods and techniques for assessing their structure and function has expanded (Chen and Kim, 2024; Graf and Kretz, 2020). At present, there are over 90,000 transcripts derived from 13,500 labeled ncRNA genes (Amaral et al., 2011; Fritah et al., 2014; Pinkney et al., 2020; Zhao et al., 2021). Catalogs of lncRNAs have been compiled in databases like GeneCaRNA, NONCODE, LNCipedia and TANRIC (Li et al., 2015) are based on transcriptional evidence. Despite the difficulties in assigning specific functions to most lncRNAs, experimental support databases that map lncRNAs to diseases are emerging (e.g., LncTarD 2.0 (Zhao et al., 2023), lncRNAWiki 2.0 (Liu et al., 2022), and lncRNAFunc (Yang et al., 2022)).

Depending on their specific mode of action, lncRNAs can act as signals, decoys, scaffolds, guides, enhancer RNAs, or even code for short peptides. Moreover, it has become evident that some lncRNAs are induced in response to environmental or cellular signals to control gene transcription. The role of lncRNAs in epigenetic changes, enhancer activity, and mRNA processing makes them attractive candidates for biomarkers and therapeutic targets (Huarte, 2015). LncRNAs’ involvement in cancer has been highlighted, and associations with tumor development, invasion, and metastasis have been proposed (Ahadi, 2021). Selected lncRNA resources focused on cancer, such as Lnc2Cancer V.2.0 (Gao et al., 2019) and Cancer LncRNome Atlas (Yan et al., 2015), utilized somatic analysis from cancer samples, experimental results, and manual curation. However, their utility in clinical applications has remained minimal.

The Cancer Genome Atlas (TCGA) has enabled large-scale profiling of lncRNA expression from tumors and normal samples from over 10,000 cancer-affected individuals, covering 33 cancer types (Tomczak et al., 2015b). This dataset provides a valuable platform for identifying lncRNAs that are selectively expressed at specific stages of cancer progression. Several lncRNAs with relatively high expression levels have been implicated in tumorigenesis. For example, HOTAIR (HOX transcript antisense RNA) interacts with the PRC2 complex to silence tumor suppressor genes. It has been shown that HOTAIR is associated with metastatic progression in breast and liver cancers (Botti et al., 2019). Similarly, MALAT1 (Metastasis Associated Lung Adenocarcinoma Transcript 1), widely studied for its role in regulating alternative splicing and metastasis, shows elevated expression in early-stage lung adenocarcinomas and is associated with poor prognosis (Kim et al., 2018) and treatment resistance (Hou et al., 2023). GAS5 has been found to act as a ceRNA for the tumor suppressor PTEN by binding to miR-21 and miR-222 in numerous cancer types (Pickard and Williams, 2015; Yang et al., 2020).

Among these abundant lncRNAs, their impact on cell identity, development, stemness, and metastasis has been reported (Li et al., 2016).

Differential expression analysis and clustering algorithms from RNA-seq data have allowed a focus on lncRNAs whose expression levels varied across tumor stages (I-IV) and thus may be utilized as biomarkers for prognosis (Ali et al., 2018; Meng et al., 2020). For example, analysis of colorectal adenocarcinoma data revealed that lncRNA CCAT1 is significantly overexpressed in stage III and IV tumors compared to early-stage cancers and adjacent healthy tissues (Nissan et al., 2012). In hepatocellular carcinoma, lncRNA DANCR was shown to promote progression from stage II to III by enhancing cancer stem cell properties (Yuan et al., 2016). In breast cancer (BRCA), lncRNA LINC00978 has been identified as upregulated in higher tumor stages and correlates with lymph node involvement and distant metastasis (Deng et al., 2016). High expression of LINC00312 distinguishes stage IV tumors in lung carcinoma and is associated with poor overall survival. In contrast, the same lncRNA in nasopharyngeal carcinoma (NPC) shows that it is negatively correlated with tumor size but positively correlated with lymph node metastasis (Zhang et al., 2013). In ovarian cancer, overwhelming evidence demonstrates that lncRNAs are essential to the pathophysiological processes, while emphasizing the lack of simple rules in the overall progression (Hu et al., 2024). These examples underscore the role of specific lncRNAs as dynamic molecular indicators of cancer stage and progression. Despite these studies, no general principles were reported for lncRNAs that dictate the cell states, tissue specificity and changes along tumor evolution. Consequently, the clinical utility of lncRNA is minimal, and their role in cancer progression remains fragmented (Mattick et al., 2023).

In our study, we created a systematic protocol for generalize a view on lncRNAs from clinical and mechanistic perspectives. The focus of our study is the identification of lncRNA signature across early and late stages of cancer progression, including the transition to a metastatic phase. Our study unified RNA-seq and clinical data from TCGA. A comprehensive, patient-centric analysis enabled a systematic analysis of lncRNA expression patterns across diverse tumor types and clinical stages, offering robust statistical power for identifying stage-associated lncRNAs. We propose an unbiased approach to reduce the collection of reported lncRNAs from 95k lncRNA genes to a reproducible set of TCGA-mapped ∼12,650 lncRNA genes that are associated with primary cancer stages and the metastatic state across major (17) cancer types. We identified a few dozen of the most informative lncRNAs that dominate any of the 7 analyzed tumor transitions and showed that the vast majority of them were overlooked. Understanding patterns of cancer progression is critical for improving clinical diagnostics and tailoring stage-specific therapies.

## Results

### Harmonization lncRNA collection for cancer research

Long non-coding RNAs (lncRNAs) are broadly defined as non-coding transcripts longer than 200 nucleotides. Over 230,000 transcripts associated with lncRNAs have been cataloged, many of which are primate-specific (e.g., compiled in LncExpDB (Li et al., 2021)). The identity of lncRNAs and the accepted nomenclature have not reached a consensus and remain largely unstable. To maintain the annotations associated with many of the reported transcripts, we applied a mapping protocol from GeneCaRNA that unified lncRNA annotation resources (see Methods).

**Fig. 1A** shows a compilation of gene compositions according to GENCODE and the human genome GRCh38.p14 (2/2022). Out of all gene types, 46% belong to lncRNAs and are assigned to their genomic locations. While lncRNAs are mostly transcribed by RNApolII (similar to coding mRNA transcripts), the remaining noncoding RNAs (ncRNAs) are primarily transcribed by RNApolI or RNApolIII and are classified as small ncRNAs (sncRNAs). Based on the GeneCaRNA mapping (Barshir et al., 2021) and filtration for contradicting mapping, the data converged to 13,257 lncRNA genes. **Fig. 1B** shows the partitioning of lncRNAs by their genomic position and transcriptional nature. We observed that most lncRNAs are intergenic (lincRNAs), intronic, and 28% of them are antisense (AS), defined by their transcriptional direction relative to the overlapping positions with coding genes. The rest of the lncRNAs display a shared promoter (i.e., bidirectional), overlap with the same directionality (sense), and more. The position of lncRNAs in the genome is a critical feature that dictates their function in cis or trans regulation. **Fig. 1C** analyzes the features of the analyzed lncRNA gene list by their transcript exon counts. The majority of the lncRNAs have ≤2 exons, and less than 1% have 10 exons. Recall that for human coding genes, the average number of exons is 9-10. **Fig. 1D** shows that the length of lncRNA transcripts is also very different from that of coding genes, with 47% being <1,000 nucleotides (nt) and only 1.1% longer than 10k nt (**Fig. 1D**). For coding genes, the average transcript length ranges from 2,000-3,000 nt, but the variability is much higher compared to lncRNA transcript length.

**Figure 1.**
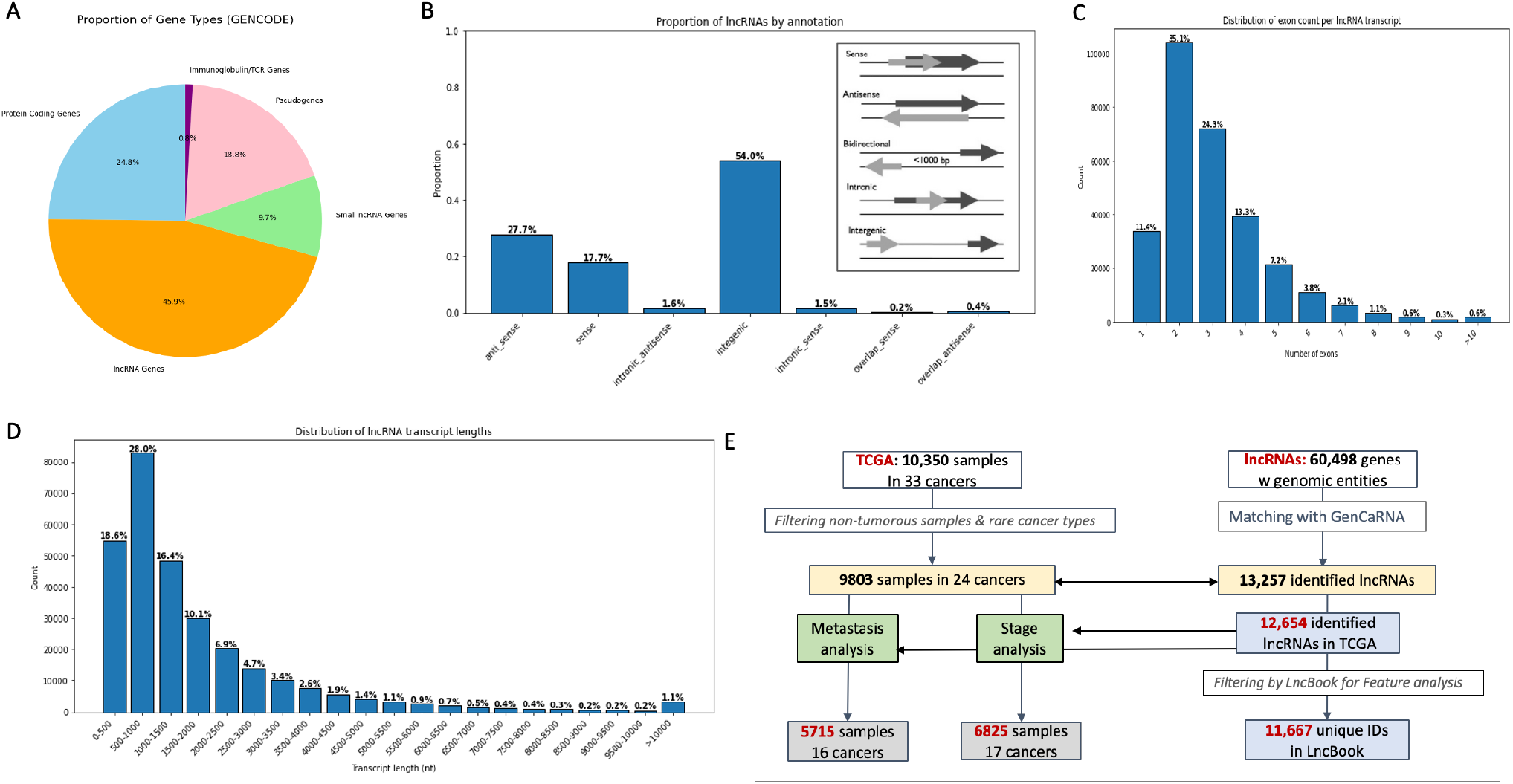
Statistics of the lncRNAs’ collection and the mapping across TCGA expression and clinical data. **(A)** Partition of the major gene types according to GENCODE Release 48 (4/2025). **(B)** The proportion of lncRNAs according to their position and transcription directionality (see inset). The intergenic lncRNA (lincRNAs) account for 54% of all links and 28% are annotated as antisense (AS). **(C)** Distribution of the exon counts per lncRNA transcripts. **(D)** Distribution of the lncRNA length by predetermined bins. For 1.1% of all analyzed lncRNAs, the length of the lncRNA transcripts is >10,000 nt.

**Fig. 1E** summarize the design of this study. We processed the TCGA public data (left) and lncRNA-mapped genes (right). The scheme emphasizes the flow for combining both resources to perform two main sets of analyses: Metastatic-based and Stage-based. The scheme used the RNA-seq and the clinical annotations modalities for each individual and each sample. Genome coordinates was used for confirming unique mapping, resulting in a final gene list of 12,654 genes. Functional and feature assignment were extracted from the lncBook dataset (Ma et al., 2019). Ambiguous mappings were removed, and an in-house cross-ID pipeline was applied (see Methods). These constraints converged to 11,667 available lncRNAs used for feature-based analysis. As each sample is labeled by stage (I-IV) and by metastatic status (M0, M1), we defined the partition of samples associated with each of the two major analyses that cover 16-17 main cancer types. We applied a conversion scheme (using the protocol from the GeneCaRNA collection, see Methods) and adhered to the common gene names and ENSEMBL gene symbols.

### Classification of major cancer types by lncRNA expression profiles

Our goal was to exhaustively study the properties of lncRNAs concerning their role in cancer progression. We focused on TCGA after filtering out rare cancer types and lncRNAs that were either not expressed at all or conflicted due to their genomic position. There were 5,715 samples across 16 cancer types and 6,826 across 17 cancer types that met the clinical annotation regarding their T-stage and M-metastatic state, respectively (**Fig. 1E**). We only considered clinical labels of these samples by T-stage (I to IV). We also combined stages I and II versus stages III and IV (primary, labeled as M0 for non-metastatic tumors) and categorized these samples as Early (E) and Late (L).

We asked whether the presence of expressed lncRNAs as the sole information from isolated samples can signify cancer types with high precision. We performed clustering by reducing dimensionality using t-SNE (see Methods) with all lncRNAs reported for each sample according to this clinical T-stage annotation (**Fig. 2A**). Each data point represents different samples, and the colored clusters were organized by the cancer types recorded for each sample from the primary tumors.

**Figure 2.**
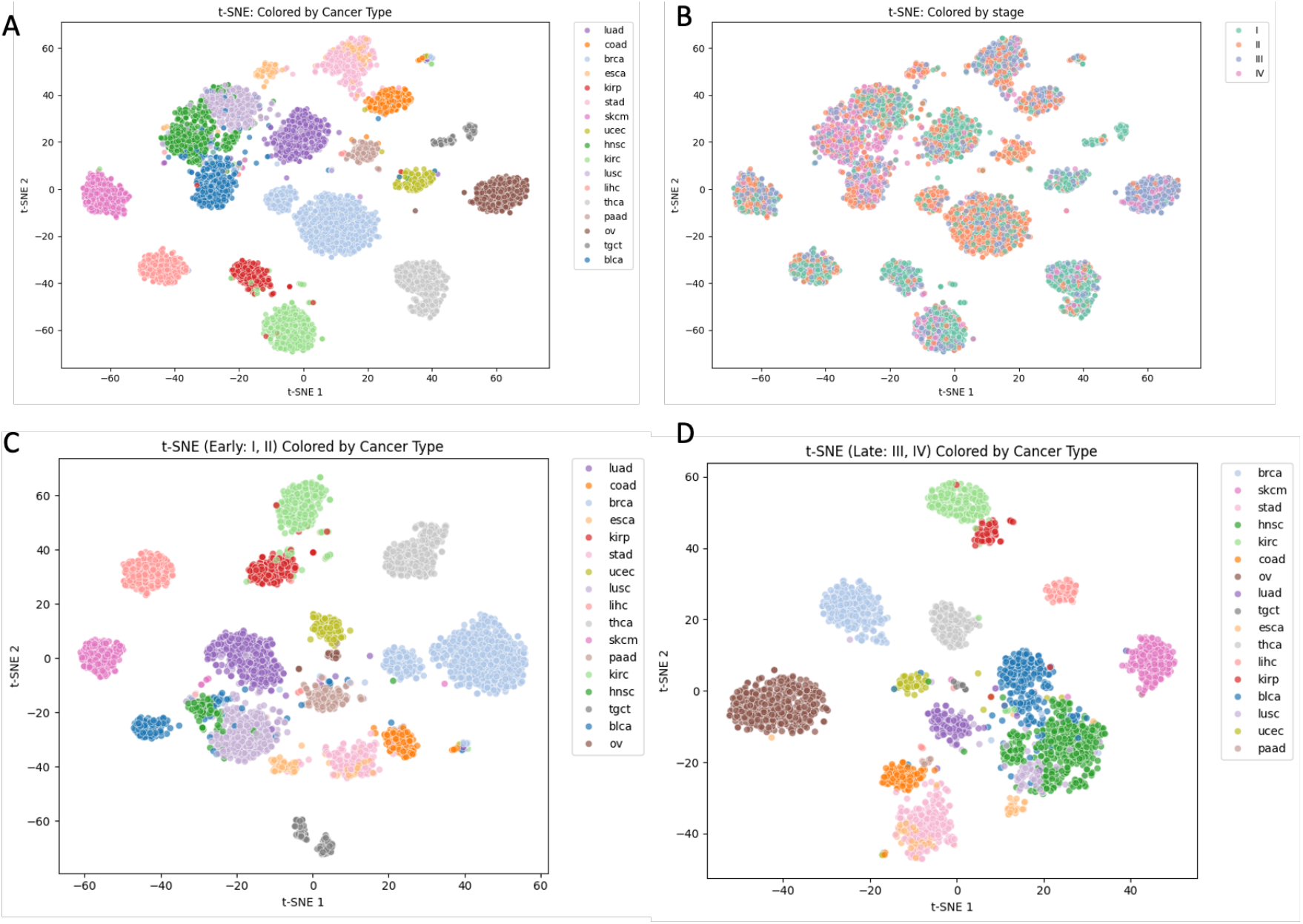
Cancer type clustering by t-SNE by lncRNAs expression profiles. The color coded for the 2D-embedding is shown for the cancer types (total 16) and for the T-stages (total 4). **(A)** The expression data used included all lncRNAs that were expressed and normalized in the analyzed samples. The resulted clusters are colored by cancer types as indicated in TCGA for each sample. **(B)**. Same 2D embedding from t-SNE as in (A). However, the color labels are according to the T-stages. **(C)** The expression data used included lncRNAs that were expressed in samples of the E-stage. Clusters are colored according to the 16 cancer types by the cancer types of the analyzed samples. **(D)** The expression data used included lncRNAs that were expressed in samples of the L-stage. Clusters are colored according to the 16 cancer types by the cancer types of the analyzed samples. Note that PADD was missing in the E-samples. Note that sample sizes differ substantially between the E and L samples. For example, while there were only a few OV samples in the E stage, the majority of the OV samples are labeled as L stage.

The t-SNE analysis validated that the lncRNAs capture information that supports a successful classification of all major cancer types. When the same analysis was performed using lncRNA expression from GTEx data of healthy tissues, a separation of all major tissues was less evident, with many of the tissues displayed as mixtures (Supplemental **Fig. S1**). We concluded that the lncRNAs collected by stages (T-stages I-IV) are informative in the dimensionality reduction performed by the t-SNE approach. We tested the possibility that the successful partition of the cancer types is dominated by their T-stage. To this end, we used the 2D visualization from the t-SNE and colored it with four colors matching T-stages I to IV (**Fig. 2B**). While it is evident that some preferences were observed for specific cancer types (e.g., T-stage II dominating KIRC), for most cancer types, the mixing of all T-stages confirmed that it is not the stage data that supports the cancer type classification.

To test whether the information from lncRNA expression specifying cancer types reflects specific T-stages, we repeated the t-SNE analysis using the partition to Early (E) and Late (L) stages. **Fig. 2C** shows that for cancers at the E stage, the partition of clusters is extremely accurate with minimal mixing. Still, a few samples were misclassified and positioned remotely from their main cluster. For example, the KIRP samples (red) slightly mixed with KIRC samples (light green). For the t-SNE representation of samples at the L stage (**Fig. 2D**), we observed a clear partition of cancer types. However, more clusters became intermixed, such as LIHC with ESCA, HNSC with BLCA, and LUSC. We concluded that the expression level of lncRNAs carries fundamental information for cancer classification, and high precision is achieved using either E or L samples for classification. Supplemental **Fig. S2** shows the results of t-SNE using lncRNA expression according to the four T-stages as non-overlapping clinical samples.

### Differentially expressed transitional lncRNAs (Tr-lncRNAs) capture cancer progression

The clinical data for T-stage and M-state are static, and each cancer type is represented by a different total number of samples and a cancer-type partition of stages and metastatic states. Supplemental **Fig. S3** shows the number of samples associated with any of the T-stages in each cancer type. For example, while there are only a few samples of stage II in OV, over 600 samples are associated with this stage in BRCA. Similarly, HSSC is dominated by stage IV (over 300 samples), but only a few samples in this stage were detected in LUSC. The Analysis of the M-state also emphasized the different characteristics of each of the 16 analyzed cancer types. KIRC is dominated by the small M0 state, while almost all PRAD tumors were classified as M0_large. To gain insight into the contribution of lncRNAs to clinical transitions, we focused on a restricted set of lncRNAs that are statistically significant across all clinical transitions. We defined a transition as a change in expression levels between consecutive states.

We used the following definitions throughout this study. For primary tumors (labeled M0) are categorized by four T-stages: stage I, stage II, stage III, and stage IV (we do not consider subdivisions into subclasses such as stage IIa and stage IIb). For the M-stage (metastatic), we categorize the clinical labels of the samples into three main categories: (i) M0_small (tumor size <2 cm), (ii) M0_large (tumor size >2 cm), and (iii) M1_large, which displays advanced large size with a metastatic state. Approximately 20 samples (across all cancer types) contradicted this classification (i.e., labeled M1_small) were eliminated and not analyzed further.

The lncRNAs that were called transitional lncRNAs (Tr-lncRNAs) specified by the following criteria: (i) A significant differential expression level (measured by fold change; log2(FC)) between consecutive states, with statistical z-value threshold of >|3|. The expression distribution by z-scores showed that using z-score >|3| accounts for ∼0.015% at each side of the distribution. For each lncRNA and each cancer type and stage, ≥30% of the samples must have a minimal level of RSEM (≥1). Example for such distribution is illustrated for BRCA (transition I ➔ II, Supplemental **Fig. S4**). For the rest of the analysis, only tails of the distribution were further analyzed. (ii) We also requested a predetermined level of coherence across samples to cope with cancer heterogeneity. Overall, we identified 7 transitions in our data, referred to as a-g transitions: (a) from stage I to stage II, (b) from stage II to stage III, (c) from stage III to stage IV. For transition (d), stage I and stage II were combined and named early (E), while stage III and stage IV are unified and named late (L). For transition of the M-state: (e) from M0_small to M0_large, (f) from M0_large to M1_large. For a more general view, we also consider transition from M0 (any tumor size) to M1. For consistency we labeled this transition ME to ML (g). Tr-lncRNAs are those that were detected in any of the 7 types of transitions. For the rest of the analyses, we considered Tr-lncRNAs across cancer types, by transition (a to g) and by the expression trend directionality in any transition. The increased and decreased expression in Tr-lncRNAs is identified by the sign of the z-scores that is positive or negative, respectively.

For simplicity, we illustrate our method of selecting Tr-lncRNAs from thousands of detected lncRNAs for a single transition of T-stage E ➔ L (transition d; **Fig. 3A**). The difference in the scale of the overall expressed lncRNAs (x-axis) and the number of TR-lncRNAs spans 2 to 3 orders of magnitude. From initial collection of 2,500-4,500 lncRNAs, only 20-80 TR-lncRNA were identified per any specific cancer type (**Fig. 3A**). When the entire collection of Tr-lncRNAs for T-stage were compiled together (transitions a-d), each cancer type is associated with ∼220 Tr-lncRNA on average (**Fig. 3B**). The number of cancer-unique was reduced to ∼60 for any of a-d transitions (**Fig. 3C**). We conclude that while still few hundreds differentially expressed lncRNAs are assigned to the T-stage transitions (transitions a-d), there are only tens of candidate that are cancer-unique. Replicating the analysis for M-state transitions (transitions e to g), resulted in similar observations, where the selected Tr-lncRNAs for M-state were reduced to only 10-80 candidates according to the expression directionality (**Fig. 3D**). The total number of Tr-lncRNAs per cancer type for any M-transitions is ∼160 in average **(Fig. 3E)**, and was reduced to only 50 for the cancer-unique subset **(Fig. 3F)**. We concluded that the use of z-score >|3| combined with a minimal fold-change in expression, apparently cancelled the large variability among cancer types that reflected the substantial difference in the number of samples labeled by the T-stage or M-state (as shown in Supplemental **Fig. S3**).

**Figure 3.**
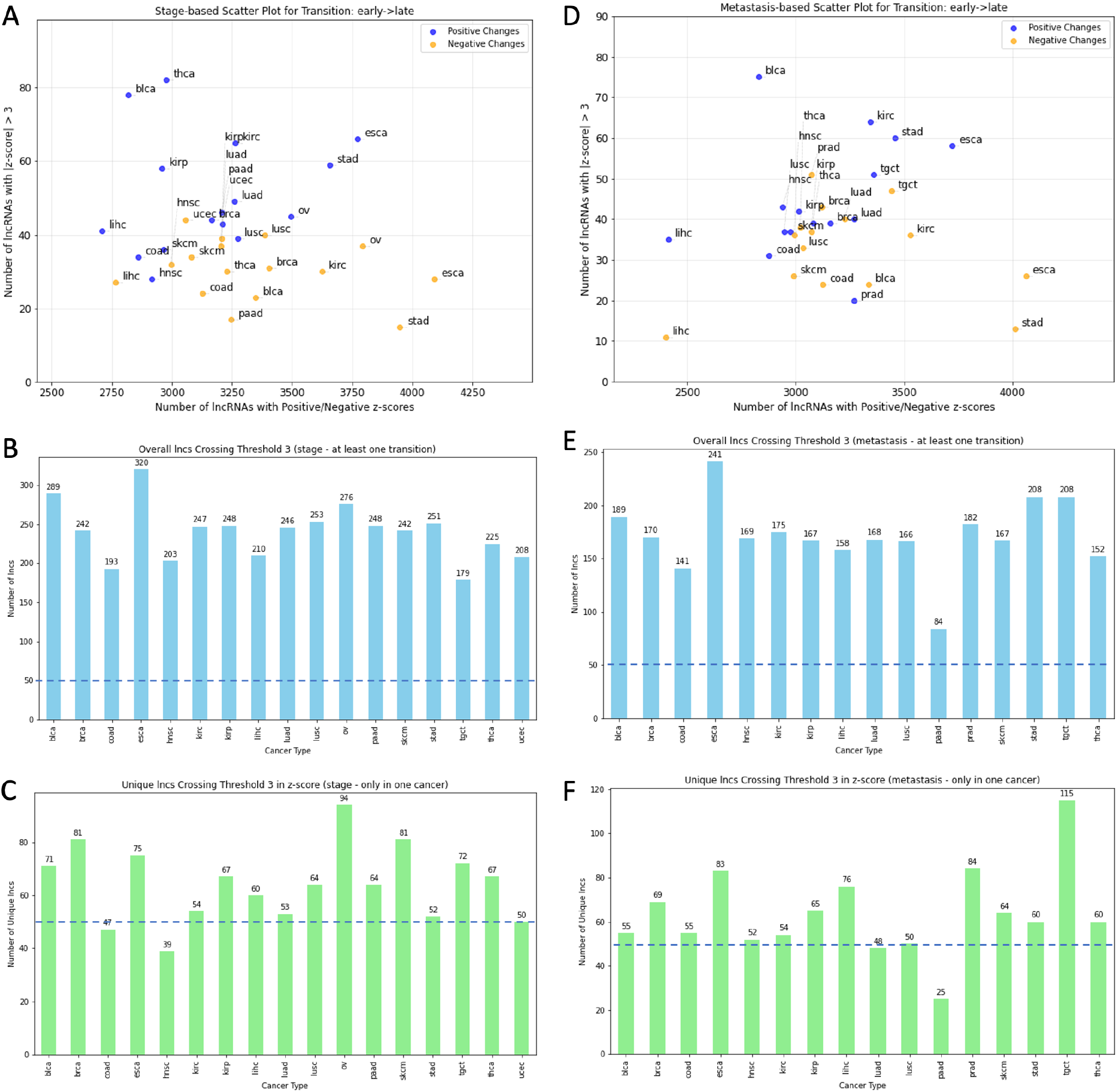
Statistics and characterization of Tr-lncRNAs by cancer types **(A)** Analysis of the Tr-lncRNAs for transition of E to L stage (transition d). The x-axis indicates the number of lncRNAs with expression level in each of the depicted 17 cancer types. The y-axis counts the number of Tr-lncRNA with partitioned to positive change (blue) and negative change (orange) that met the z-value threshold (z values >|3|. **(B)** Summary for all Tr-lncRNAs that were reported with at least one T-stage transition (transitions a-d) across 17 cancer types. The dashed line was added for a visualization reference. **(C)** A subset of Tr-lncRNAs from (B) that were only reported within a single cancer (cancer-unique) is shown. The dashed line was added for a visualization reference. **(D)** Analysis of the Tr-lncRNAs for transition of metastatic ME to ML state (transition g). The x-axis indicates the number of lncRNAs with expression level in each of the depicted 16 cancer types. The y-axis counts the number of Tr-lncRNA with partitioned to positive change (blue) and negative change (orange) that met the z-value threshold (z values >|3|. **(D)** Summary for all Tr-lncRNAs that were reported with at least one M-stage transition (transitions e-g) across 16 cancer types. The dashed line was added for a visualization reference. **(E)** A subset of Tr-lncRNAs from **(D)** that were only reported within a single cancer (cancer-unique) is shown. The dashed line was added for a visualization reference. For caner type abbreviations see Supplemental **Table S1**.

### Tr-lncRNAs connectivity map and network across cancer types

**Fig. 4A** is a schematic view of the main analyses for identification of Tr-lncRNAs. Each cancer type (total 17) is analyzed by seven transitions (labelled a to g). Transitions a-d are associated with T-state and e-g with M state. Tr-lncRNAs are defined by a minimal expression across a minimal number of samples (see Methods) and z-scores of >3 and <-3. The five most significant Tr-lncRNAs for each cancer types with positive and negative z-values are reported. For simplicity, we demonstrate a connectivity matrix with T-stage transition of E➔L. **Fig 4B** shows that the degree of sharing of Tr-lncRNAs is quite minimal and for most cancer type pairs, only 2-3% of the lncRNAs are shared (absolute numbers of Tr-lncRNAs are indicated in the upper right triangle, **Fig. 4B**). The maximal level of similarity is bounded by less than 10% (for the pairs of BLCA and BRCA). The Tr-lnRNA shared matrix for all other T-stage transitions (transition a-c) and M-state transitions (transition e-g) are shown in Supplemental **Fig. S5)**.

**Figure 4.**
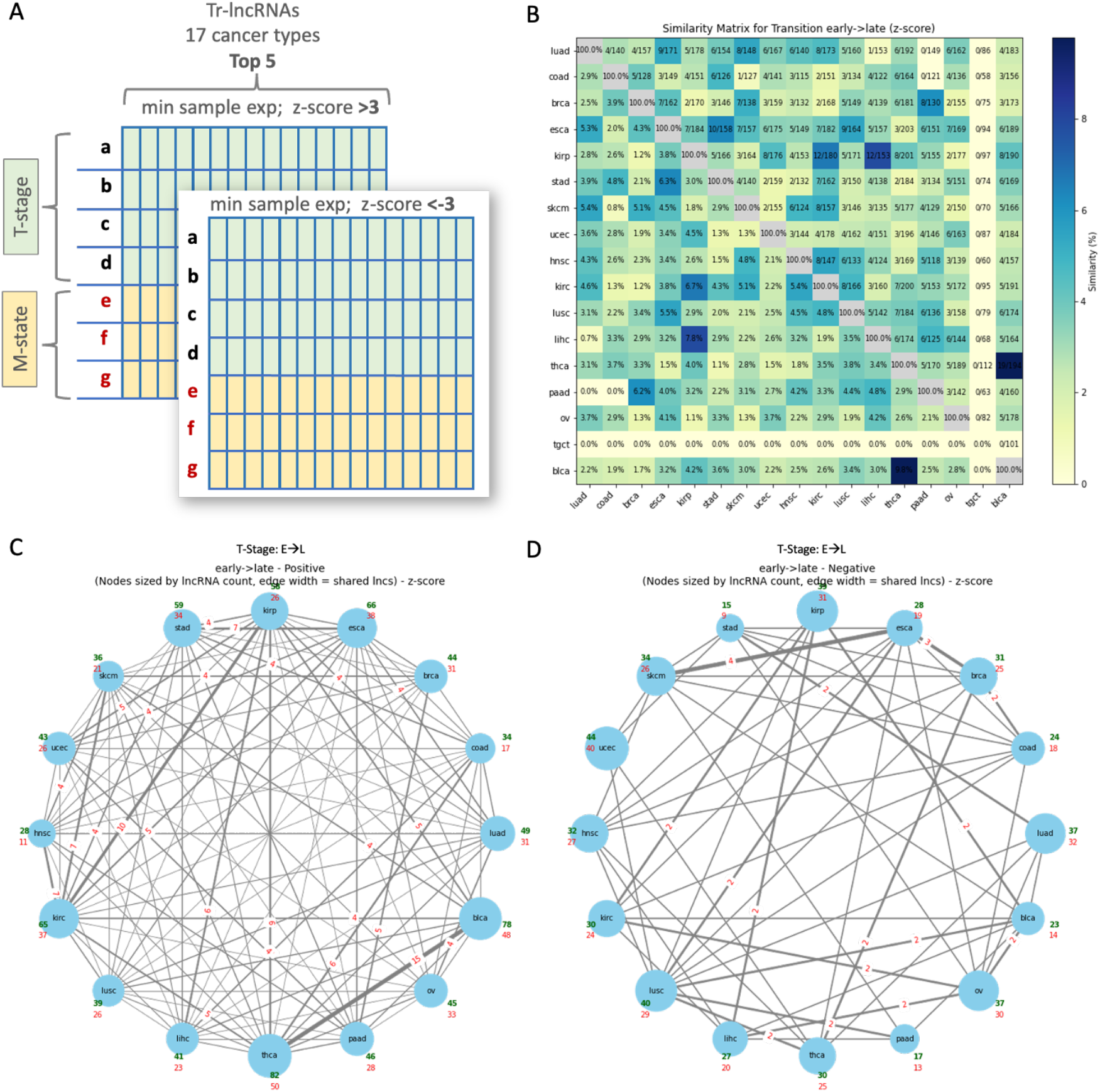
Tr-lncRNAs among cancer types. **(A)** Schematic view of the main analyses for identification of Tr-lncRNAs. Each cancer type (total 17) is analyzed for 7 transitions (labelled a to g). Transitions a-d are associated with T-state and e-g with M state. Tr-lncRNAs are defined by a minimal expression across samples and z-scores of >3 and <3. The 5 most significant Tr-lncRNAs for each cancer types with positive and negative z-values are reported. Heatmap for **(B)** Heatmap for the T-stage E to L transition (transition d) on 17 cancer types (note that tgct has not been analyzed) colored by the percentage of sharing with any other cancer types (bottom left), and in a symmetrical position above the diagonal, the actual number of shared Tr-lncRNAs from the total number of potential lncRNAs. **(C)** Connectively map of all cancer types (nodes) and the transition of T-stage E➔ L for the positive z-values. The number are indicated for ≥4 of shared Tr-lncRNAs. **(D)** Connectively map of all cancer types (nodes) and the transition of T-stage E➔ L for the negative z-values. The number are indicated for ≥2 of shared Tr-lncRNAs. Edges indicate shared Tr-lncRNA connecting two nodes (i.e., cancer types). The size of the node reflects the number of samples. and the width of the edges indicates the number of shared Tr-lncRNAs (marked on the edge). In red fonts are the number of unique Tr-lncRNAs for each cancer type.

We assessed the overlap in expression trends of Tr-lncRNAs across all cancer types. **Fig. 4C** shows the results for the T-stage E ➔ L transition for the positive z-value (i.e., upregulating as the cancer progress from E ➔ L stage). We observed that the network of Tr-lncRNAs that are characterized by a suppression in expression at the E ➔ L transition (i.e., negative z-values) is less dense and most edges associated with only a single Tr-lncRNAs (**Fig. 4D**). A similar observation was associated with most of the transitions. Nevertheless, the absolute number of such shared Tr-lncRNAs is small as reflected by the edge weigh. Furthermore, we observed that about 60% of the Tr-lncRNAs unified by positive z-scores are unique (red font in cancer types nodes; **Fig. 4C)** and the fraction of unique among the Tr-lncRNAs for the negative z-scores (**Fig. 4D**) is even higher. For example, among the 44 Tr-lncRNAs associated with USEC, 40 of them are unique. A network connectivity for a transition of ME➔ML for the positive and negative expression directionality is shown in Supplemental **Fig. S6**). We conclude that this exhaustive connectivity analysis across all 17 cancer types highlights a relatively small, coherent set of candidate lncRNAs as candidates for functional investigation.

### Pan-cancer view highlights small set of Tr-lncRNAs that specify cancer progression

Based on the high specificity of Tr-lncRNAs for each cancer type, we inspected cases in which Tr-lncRNAs dominates T-stage or M-state transitions across as many cancer types as possible. To this end, we performed a pan-cancer view in which the most influential 50 most Tr-lncRNAs are listed according to high occurrences in as many of the cancer types, and their maximal accumulated z-values (by absolute values). Supplemental **Table S2** lists Tr-lncRNAs for transition d and g (for all ≥2 cancer types). We illustrate the approach for the two transitions of M0-small to M0_large (i.e., tumor growth) and the M0_large to M1_large (i.e., switch to metastatic state).

Table 1 summarizes information for the 14 TR-lncRNAs that overlapped between transition e and f using the top 50 Tr-lncRNAs of each transition. In terms of genomic transcriptional mode, there are two intergenic genes (i.e., lincRNAs), three antisense lncRNAs of the genes: (i) HAND2, an embryonic developmental gene; (ii) ST8SIA6, a gene known in numerous cancers, with increased expression that is associated with decreased survival. (ii) DIO3OS a bidirectional positioning with DIO3 which is central to thyroid metabolism. In the context of cancer cells, the DIO3 promotes proliferation and invasion. The other lncRNAs were extensively studied include XIST and USP9Y that acts in the sex chromosomes. The common feature is that gene genes are sensitive to their level of expression for fulfill their function. The knowledge for the rest of the lncRNAs is quite sporadic. We argue that the discovery of a small set of lncRNAs that are involved in different aspects of tumorigenesis is of a clinical relevance.

Based on the list of top ranking lncRNA for M transition, we tested the coordinated expression across cancer types and the trend of expression per each of the identified lncRNAs (**Fig. 5**). To this end we show the z-score for shared 14 cancer types of representative Tr-lncRNAs. **Fig. 5A** shows the coordinated behavior of XIST across the different cancer types and the coherence between the z-values of transition e and f. The XIST shows a strong positive correlation for the two M-state transitions r=0.59 and p-value=0.02 (**Fig. 5A**, right). The role of XIST is well established and while for many cancer types it is strongly suppressed with tumor growth. For example, in lung cancers (LAUD and LUSC), the suppression is extreme (z-value ∼-15; (**Fig. 5A**). Notably, across almost all cancer types, the transition trends were consistent (i.e., positive or negative).

**Table 1.** Shared Tr-lncRNAs for the M-state based on the pan-cancer across 15 major cancer types.

| lncRNA | Chr. Location | Function summary<sup>a</sup> | TCGA Cancer Type<sup>b</sup> |
| --- | --- | --- | --- |
| AFAP1-AS1 | 4p16.1 | Onc; promotes metastasis via miRNA sponge | LUSC, OV, PAAD, ESCA, STAD, BRCA, COAD, LIHC, LUAD |
| DIO3OS | 14q32.2 | TSG; miRNA sponge | LIHC, BRCA, PAAD, THCA |
| HAND2-AS1 | 4q34.1 | TSG; inhibits proliferation, metastasis | COAD, CESC, LIHC, BRCA |
| LINC00668 | 18p11.31 | Onc; promotes proliferation, EMT | LIHC, LUAD |
| LINC01234 | 12q24.13 | Onc; promotes proliferation, migration via miRNA sponge | ESCA, KIRC, LIHC, STAD, COAD, BRCA, OV, LUAD |
| MIR205HG | 1q32.2 | Onc; ceRNA regulating tumor processes | CESC, PRAD, THCA |
| RP11-1143G9.4 | 8q24.21 | Poorly characterized | COAD |
| RP11-152H18.3 | 7q31.1 | Poorly characterized | ? |
| RP11-334J6.7 | 11p15.5 | Poorly characterized | LIHC, ? |
| RP11-498C9.13 | 11q24.1 | Poorly characterized | GBM, ? |
| RP5-940J5.9 | 17q24.3 | Upregulated in COAD; therapeutic potential | COAD |
| ST8SIA6-AS1 | 10q24.1 | Onc; promotes proliferation, invasion | LIHC |
| USP9Y | Yp11.2 | TSG; deubiquitination | LUAD |
| XIST | Xq13.2 | X-inactivation | HNSC, BRCA, CESC, PRAD |

<sup>a</sup>TSG, tumor suppressor gene; Onc, oncogene; EMT, EMT, epithelial-to-mesenchymal transition. <sup>b</sup>TCGA, abbreviation for major cancer types.

**Figure 5.**
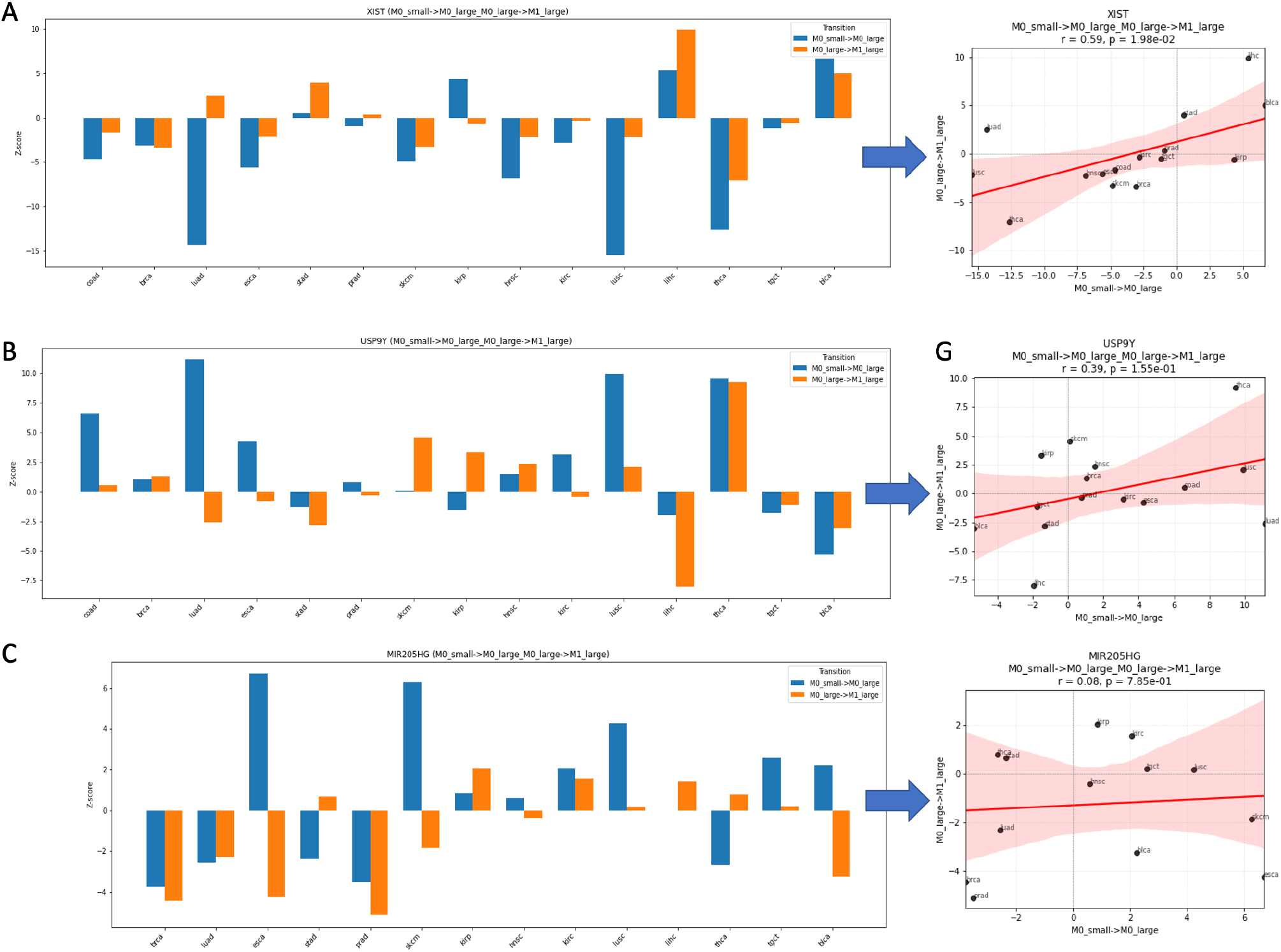
Shared Tr-lncRNAs for M-state transitions based on pan-cancer analysis across cancer types. For each cancer types the Tr-lncRNAs from pan-cancer analysis are shown with two transitions (e and f) and the associated z-values (y-axis). On the right, a scatter plot where each datapoint is a cancer type and the value associated with the two tested transition. Pearson correlation (r) is calculated along with the statistics (by p-value), with 95% confidence interval (CI) colored in pink. The analyzed lncRNAs tested are: **(A)** XIST **(B)** USP9Y **(C)** MIR205HG. Note that the y-scale is gene specific. The order of the cancer types is preserved across the different Tr-lncRNAS tested.

Another shared gene is the USP9Y that is involved in many cancer types (**Fig. 5B)**. For this lncRNA the amplitude of the gene expression is less extreme, but tend to keep consistent among cancer types (r=0.39). Inspecting XIST and USP9Y together show that in almost all cancer types, they exhibit opposite trend. For example, in THCA XIST is suppressed in both transitions while exhibits strong positive expression trend for USP9Y. The same contrasting trends is observed for most cancer types. **Fig. 5C** shows that there are other lncRNAs (e.g., MIR205HG) with no coherence between the two M-state transitions (r=0.08). In such cases, we observed inconsistency for the transition along the primary tumor growth and the establishment of metastasis.

**Fig. 6A** illustrate the pan-cancer analysis scheme for the top 50 Tr-lncRNAs identified by their occurrence in maximal number of cancer types and the maximal sum of the z-scores. Applying the pan-cancer analysis for identifying the most informative lncRNAs for T-stage for the a-c transitions confirmed the occurrence of XIST, UCA1, H19, DIO3OS, MIR205HG and USP9Y as well as two antisense lncRNAs (AFAP1-AS1 and VCAM-AS1; **Fig. 6B)**. The overlap of Tr-lncRNAs among transition a-c with the top 50 genes per each transition further emphasize that the overlap in lncRNAs is only moderate, and only 6 lncRNAs, 5 of which are poorly characterized were identified across all these transitions, supporting the notion that lncRNAs can serve as sensitive indicators for specific stages in cancer progression. Among the shared lncRNAs, RP5-940J5.9 shows maximal variation across all cancer types (Supplemental **Table S2**).

**Figure 6.**
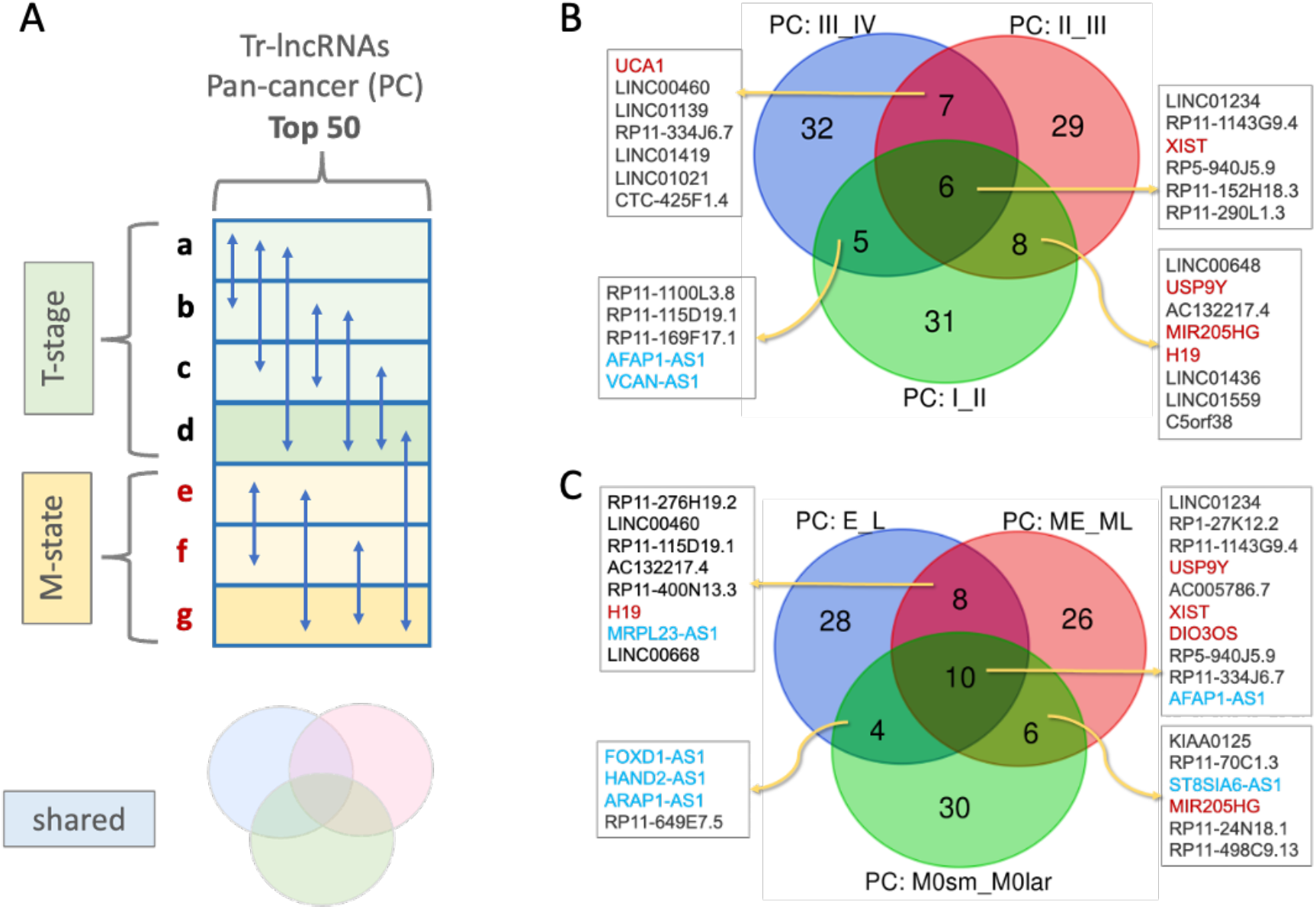
Pan cancer analysis of Tr-lncRNAs. **(A)** Scheme of all 7 transitions (a-g). The T-stage and M-state transitions are colored green and yellow, respectively. The d and g transition are indicative for the T-stage E➔ L and the M-state ME➔ ML. The vertical arrows marks comparison between any pairs of transitions. The Venn diagram symbol is used to identify the unique versus the shared Tr-lncRNAs for any combinations of transitions. **(B)** Pan cancer (PC) view of T-stage transitions a-c. The top 50 Tr-lncRNAs across the cancer types for T-stage transitions is shown by the Venn diagram. **(C)** Pan cancer (PC) view of M-state transitions e-g. The top 50 Tr-lncRNAs across the cancer types for M-state transitions is shown by the Venn diagram. Overlapping Tr-lnRNAs names are shown. Colored blue are Tr-lncRNAs that were annotations of antisense (AS). Names of previously studied lncRNAs that were assigned with accepted names are colored red (e.g., XIST, H19, UCA1).

We asked whether there are Tr-lncRNAs that are shared between T-stage and M-state transitions. To this end, we compared M-state transition that specify tumor growth (M0_small to M0_large) as well as the lower resolution transitions marked as E➔L and ME➔ML (**Fig. 6C**). We observed that the overlapping set is quite moderate (total 28 lncRNAs) with 10 that are shared between transitions d, e and g. Comparing the observation in **Fig. 6B** we show that XIST, H19, DIO3OS, MIR205HG and USP9Y are shared (not UCA1). Altogether, there were 6 antisense lncRNAs, 3 of them mark the tumor growth (transition e and d), irrespectively to the switch to metastatic state (FOXD1-AS1, HAND2-AS1 and ARAP1-AS1).

### Identifying most significant T-stage Tr-lncRNAS candidate per cancer type

We further analyzed and listed 5 top genes for each cancer type, each transition (T-stage and M-state transitions) and expression trend directionality. We focus on the results for T-state transitions a-d. We show cancer types as showcase for further generalization. **Fig. 7A** shows the results from BLCA. We observed that among the top 5 Tr-lncRNAs the z-score was very significant and many of the Tr-lncRNA were most significant than z-score >7 (**Fig. 7A**, dashed vertical line). We identified XIST among the top list in transitions a, b and d and UCA1 as the most significant in transition c. The other gene that is among the top of transition d is MIR99AHG, a host gene of miR-99a, let-7c and miR-125b-2. These miRNAs act mostly as tumor-suppressor in various cancers and involved in targeting oncogenes. The analysis performed on COAD (**Fig. 7B**) also shown significant Tr-lncRNAs. In this case, HOTAIR was identified in transitions c and d. A general observation emphasizes the occurrence of antisense lncRNAs (AS) among the top listed genes but mostly poorly studied lncRNAs (e.g., prefix PR11, ACO). **Fig. 7C** (STAD) and **Fig 7D** (PAAD) show that the top 5 Tr-lncRNAs have slightly lower z-values. Moreover, a few genes were observed across cancer types. For example, AFAP1-AS1 that was signified in the M-state **(Table 1**), also identified in STAD transition c (**Fig. 7C**) and in PAAD in transition a and d (**Fig. 7D**). We observed that for in cancer types, the majority of the identified TR-lncRNAs are novel and were overlooked. This also include lncRNAs that were identified as the most significant lncRNAs. For example, the z-score of RP11-462G2 in PAAD is 7.63, the most significant of any other listed lncRNAs in this cancer (**Fig. 7D**). While it is poorly characterized, recently it was proposed to be part of a complex regulatory network influencing tumor invasion in colorectal cancer (Zhou and Li, 2022). Similarly, inspecting the Tr-lncRNA with z values >3 identified top genes that are negatively expressed along the transitions (i.e., z value <3). For keeping the analyses concise, these are not discussed (Supplemental **Table S3**). We concluded that by using our strict thresholds of differential expression and focusing on the specific cancer type, selected candidate list of genes can be directly used for clinical characterization purposes and as cancer progression indicators.

**Figure 7.**
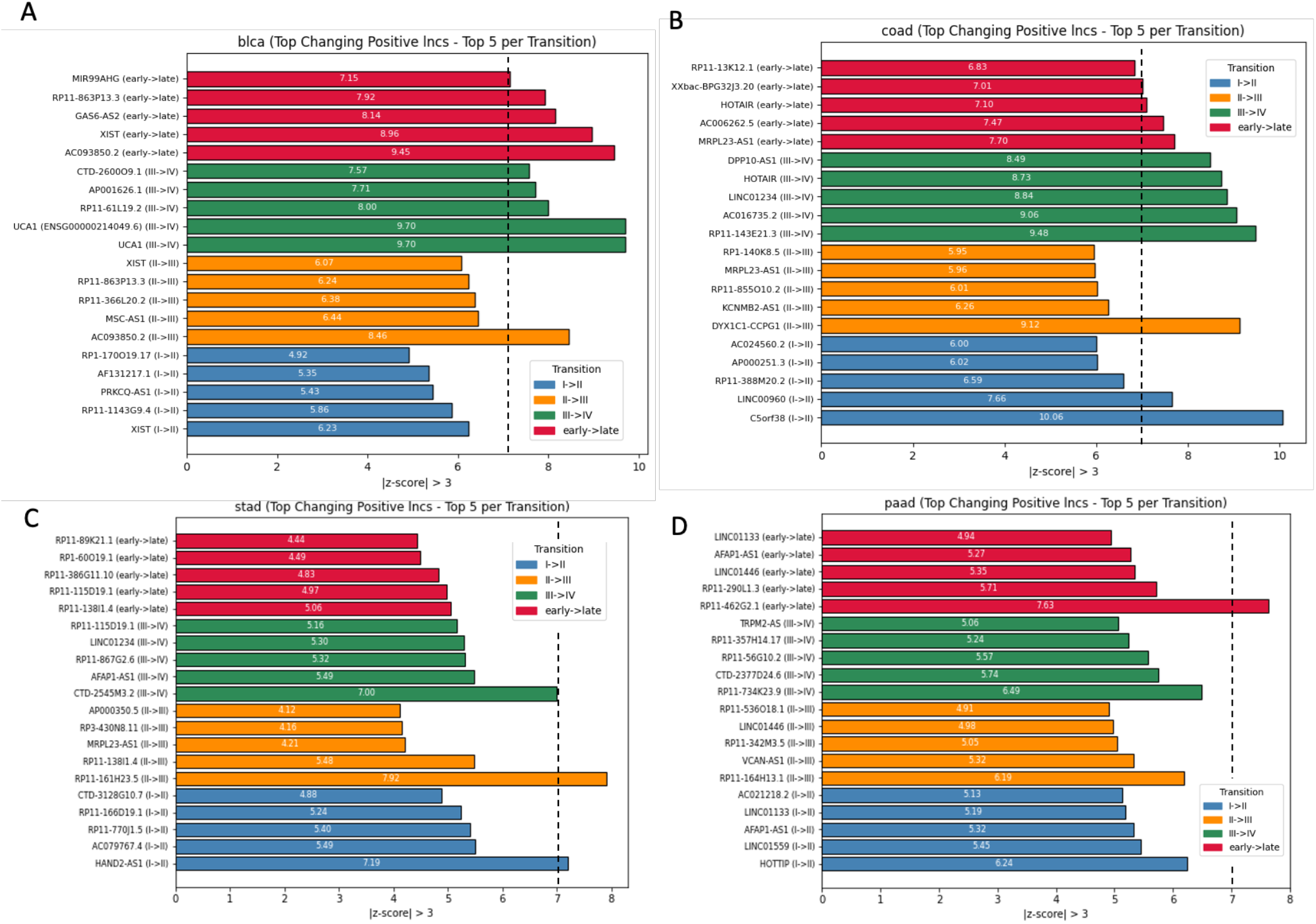
Top 5 lncRNAs per transition in 4 different cancer types with increase expression (z value >3) for all cancer transitions, marked by colored bars. **(A)** BLCA for T-stage a-d transitions. **(B)** COAD for T-stage a-d transitions. **(C)** STAD for M-state e-g transitions. **(D)** PAAD for M-state e-g transitions. The vertical dashed line marks z-score =7 for visualization purpose only. The values of the z-score for each lncRNA is indicated within the relevant bar. Note that a few lncRNAs may appear in more than one transition.

### A mechanistic view on the function of candidate Tr-lncRNAs

The identification of only a few Tr-lncRNAs associated with a specific cancer type provides an opportunity to systematically assess the possible mechanism of action. We tested the possibility that upon transition, high expressing lncRNAs may be coupled (in a negative or positive direction) with the sense coding gene at their vicinity. We illustrate a few examples to expose the general picture that emerges from the genomic location of the Tr-lncRNAs.

**Fig. 8** focuses on a few examples of such lncRNAs that are either annotated as AS or cases where inspecting the genome position suggests cis-regulatory transcription mode of action. **Fig. 8A** shows the genomic position of RP5-940J5.9, a poorly characterized lncRNA that we have identified in the pan-cancer view for M-state (**Table 1**) and also among the 4 Tr-lncRNAs shared across T-stage transitions (**Fig. 6**). The genomic position of this lncRNA confirm its position in a segment with enhancers within an extremely active region, at the vicinity of GAPDH. Actually, RP5-940J5.9 is an antisense of GAPDH. While poorly characterized, this lncRNA was observed in pediatric B-ALL (B-cell acute lymphoblastic leukemia) that were strongly upregulated relative to healthy samples (Affinito et al., 2020). **Fig. 8B** focuses on DIO3OS which supports a bidirectional transcription. As expected, a strong positive correlation was observed across tissue specifically for the coding gene of DIO3 and lncRNA DIO3OS expression (see GTEx track, **Fig. 8B**). As for the RP5-940J5.9, DIO3OS was identified by a pan-cancer view for M-state (**Table 1)** and T-stage transitions **(Fig. 6)**.

**Figure 8.**
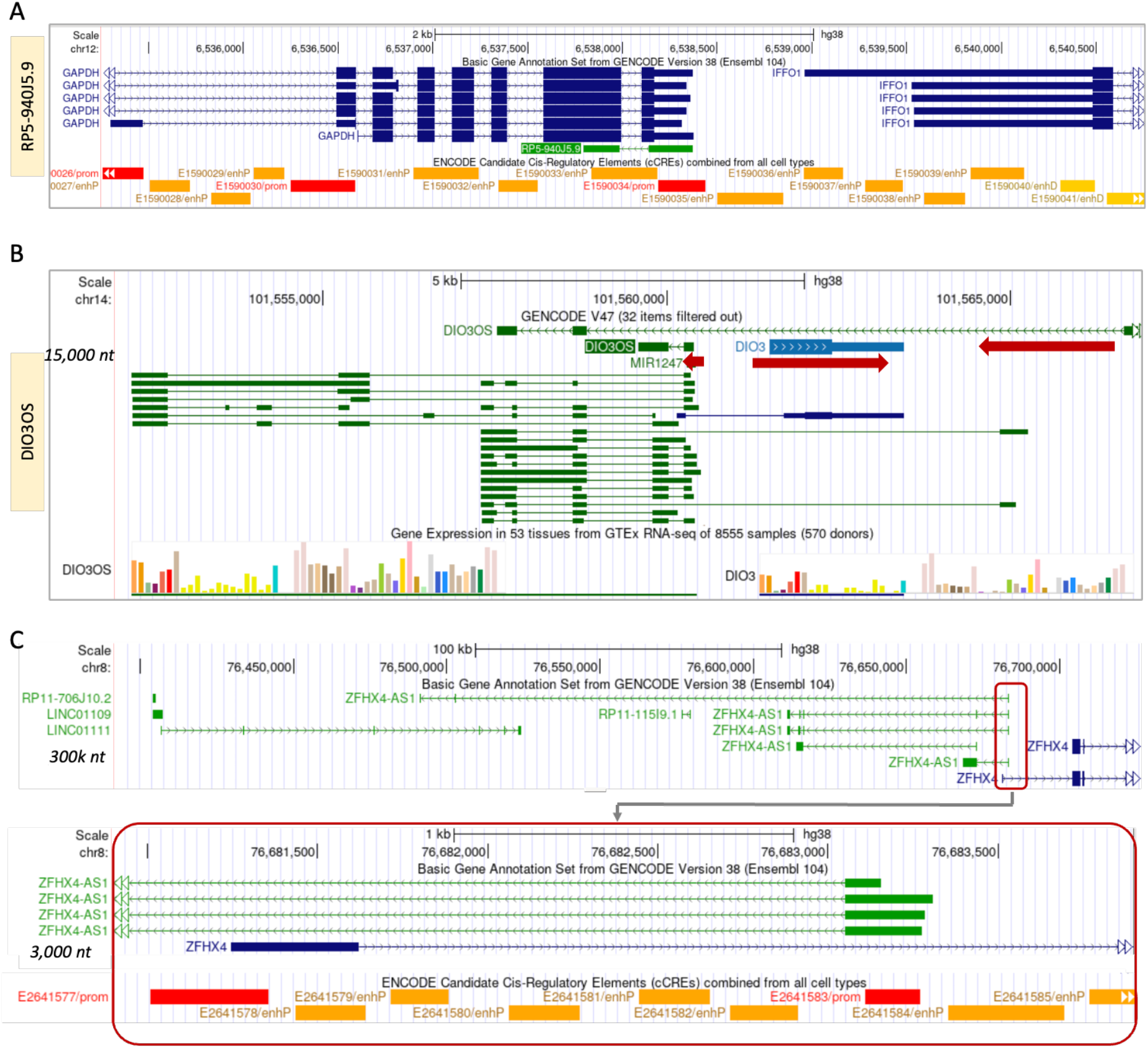
Genomic view on transcripts of Tr-lncRNAs by pan-cancer view. **(A)** The lncRNA RP5-940J5.9 is shown in the context of GAPDH and the rich promotor enhancer regulatory elements as presented by Encode. RP5-940J5 may act as AS for GPDH that is highly expressed in many cancer types. **(B)** A region of 15k nt covers the coding gene of DIO3 (+ strand), miR-1247 and DIO3OS (−strand). The multiple lncRNAs of DIO3OS are shown in green. Transcriptional direction is indicated by red arrows. The GTEx gene expression are shown with multiple tissues, expression profile of DIO3 and DIO3OS strongly correlate. **(C)** A region of 300k nt covers a segment rich with lncRNAs (in green) and several transcripts of ZFHX4-AS1. Zoom in to a subregion of 3000 nt shows AS transcriptional position relative to the ZFHX4. The ZFHX4-AS1 is positioned in promoter-enhancer rich segment.

**Fig. 8C** highlight the genomic context of the Tr-lncRNA ZFHX4-AS1 which was significant for T-stage of E➔L transition. It is an exceptionally long lncRNA, derived from 8 exons that spans ∼300k nt in Chr8 (8q21.13). This lncRNA is predominantly expresses in brain and testis. It has been implicated in a range of processes including cell proliferation, metabolism, and immune cell infiltration. Zoon in of this region (3,000 nt, **Fig. 8C**) highlights its relevance to promoter-enhancer regulation (enhancer RNA, eRNA; **Fig. 8C**, bottom panel). It was shown that in ovarian cancer (OC) from TCGA, the expression of ZFHX4-AS1 is associated with poor prognosis and was proposed as an independent prognostic factor for OC survival. Moreover, the expression of ZFHX4-AS1 is linked to pathways like TGF-beta and PI3K-Akt signaling and signifies advanced disease stages and immune cell infiltration (Zong et al., 2022). it is likely that the region that is packed with enhancers-promoters supports the expression of ZFHX4 (e.g., in glioblastoma). In several cancers, the ZFHX4 regulate epithelial-mesenchymal transition (EMT) and reprogramming the extracellular matrix (ECM) to promote migration.

### Tr-lncRNA molecular features are significantly different compared to the lncRNA collection

The selection of Tr-lncRNAs ignored all molecular features while only considering expression levels and a threshold on sample consistency. We used a rich collection of annotations to assess statistical measures that are enriched or underrepresented in the unified set of Tr-lncRNAs (a total of 2,399 unique lncRNAs) relative to all lncRNAs that were mapped to TCGA (Fig. 1E) and annotated according to lncBook (Li et al., 2023). Information on transcript expression, protein capacity, genomic features (e.g., number of exons, exon length, gene length), conservation (e.g., conservation along the phylogenetic tree), and subcellular annotations yielded 143 features. We have not included features on lncRNA interactions, methylation, or genomic variants. Comparing the statistics of the background set (all lncRNAs) with Tr-lncRNAs identified 40 (out of 143) features that met the FDR p-value <0.05 (**Fig. 9A**). The fraction of small proteins is higher in Tr-lncRNA relative to the background collection. **Fig. 9B** shows that Tr-lncRNAs are overall longer, with more exons and longer exons. Other transcript features show that the number of transcripts per gene is substantially higher than the background of all lncRNAs, and this also applies to poly A sites (**Fig. 9C**). The 2-fold higher fraction of small proteins among Tr-lncRNAs relative to other lncRNAs strongly suggests that this set may be associated with high fraction of encoded protein and peptides (**Fig. 9D**). Interestingly, none of the 40 subcellular annotations were significant, and only a very few of the many conservation annotations (totaling 30) met the FDR threshold. The list and statistics for all 143 features are included in Supplemental **Table S4**.

**Figure 9.**
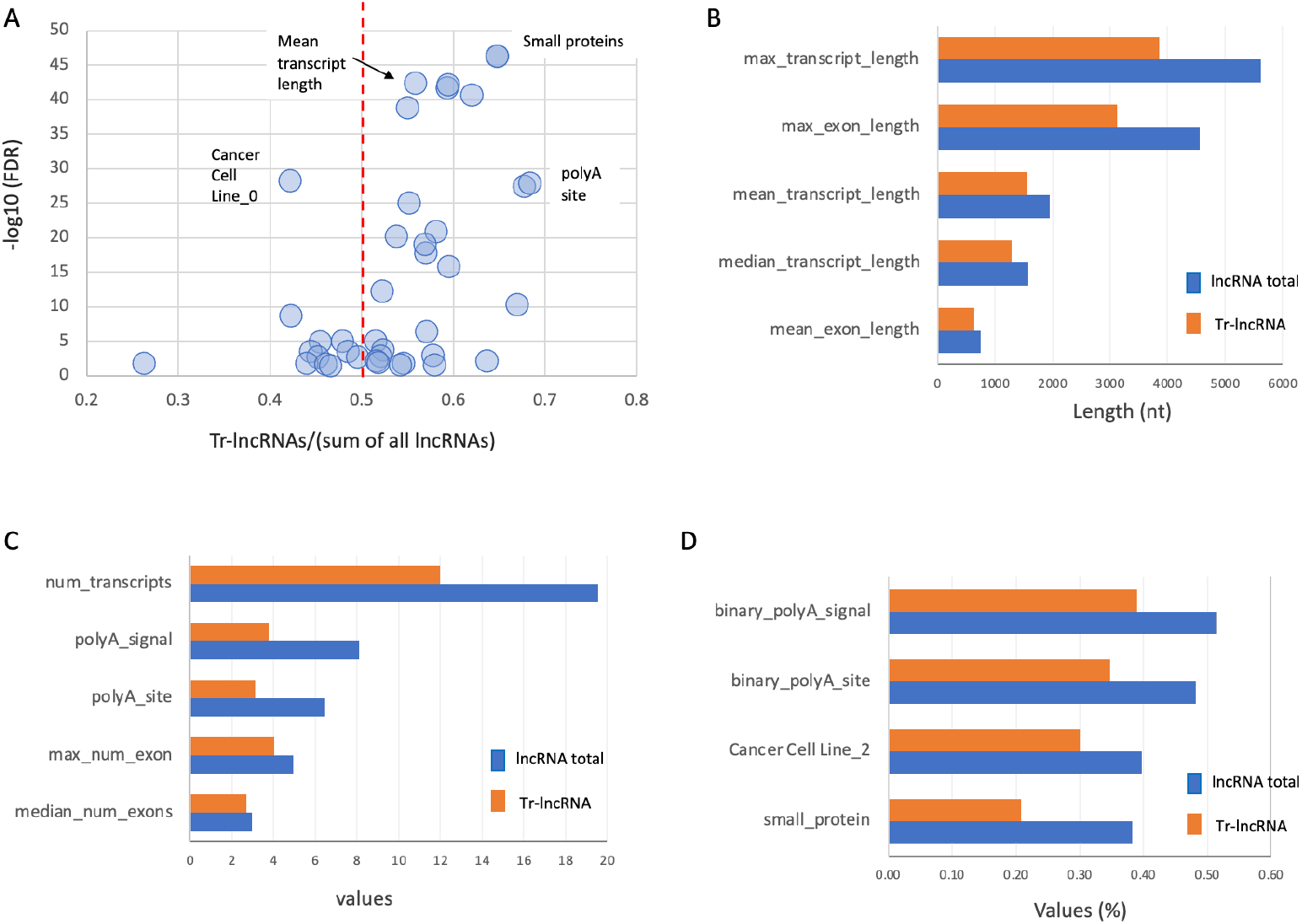
Feature enrichment of Tr-lncRNAs based on annotations extracted from the lncBook collection on genomics, transcripts, conservation, and protein capacity (see Methods). **(A)** Scatter plot of the 40 significant features. The ratio of Tr-lncRNA values to the sum of all values (Tr-lncRNA and all lncRNAs) is shown. The vertical line (0.5) indicates an identical value for both lncRNA groups. Values on the x-axis >0.5 signify enrichment in Tr-lncRNAs, and values < 0.5 are underrepresented in the Tr-lncRNA group. For example, a 2-fold enrichment is translated to value >0.67. **(B)** Values for features associated with transcript length. **(C)** Features associated with distinct characteristics and occurrences. **(D)** Binary and categorical features and their prevalence. All 143 features and their statistics are available in Supplemental **Table S4**.

## DISCUSSION

In this study, we present a comprehensive harmonization of lncRNA data relevant to cancer. Our curated reference set of approximately 12,500 lncRNAs provides a high-confidence collection, with the majority supported by multiple catalogs and annotation platforms such as lncBook (Ma et al., 2019), NONCODE (Fang et al., 2018), and GENCODE (Frankish et al., 2023). An effort to characterize the landscape of lncRNAs in 2015 confirmed that 80% of them lack any annotation (Iyer et al., 2015). By integrating resources with multiple annotations and aligning lncRNA identifiers across these resources, we achieved consistent indexing that facilitates downstream analysis and cross-database utilization. Matching the lncRNAs to the transcripts reported by TCGA allowed to assess the contribution of lncRNAs to the major phases of cancer progression. Our approach ensures that we maintain robust and statistically solid discovery. Our protocol for identifying the most informative lncRNAs in cancer transitions identified an unprecedented large number of novel lncRNAs, most of them lack supporting evidence for their potential function.

The collection of all candidate list includes 2,399 Tr-lncRNAs that stand out due to their fold changes across cancer transitions (i.e., with 5 most extreme positive and 5 most negative z-scores, across 7 transitions and across 16 cancer types; Fig. 7). Among these Tr-lncRNA candidates, we successfully recovered many well-characterized functional lncRNAs, including H19, MALAT1, XIST, UCA1, HOTAIR, USP9Y, and others. Notably, the statistical properties of known and novel lncRNAs showed no significant differences. All candidates were filtered using stringent criteria to reduce noise. Our approach integrated statistical rigor (z-score >|3|) with biological expression coherence across samples, supporting robust expression patterns suggestive of functional relevance (Liu et al., 2022). The result is a ranked list of overlooked lncRNAs with strong potential relevance to cancer progression (Liu et al., 2017; Pandey and Kanduri, 2022).

It has been suggested that lncRNA expression is sufficient to robustly classify major cancer types (Iyer et al., 2015). We observed the ability to classify cancer types by lncRNA expression alone with greater success compared to the classification of healthy tissues from GTEx by lncRNA expression profiles. Cancer heterogeneity and multiple clonality in tumors make classification by mRNA profiles challenging in bulk analysis (Capper et al., 2018). In contrast, healthy tissues (e.g., GTEx data) were successfully classified using unsupervised methods. This discrepancy was attributed to the robust and reproducible transcriptional networks of mature tissues (Saha et al., 2017). A similar observation was associated with cancerous and their normal tissue counterparts using miRNA profiling (Rasnic et al., 2017). These findings align with the emerging view accordingly the lncRNAs reflect cancer-specific transcriptional networks with high sensitivity and specificity. To capture the dynamic regulation, we introduce the concept of transitional lncRNAs (Tr-lncRNAs), defined by coherent expression shifts across clinical stages (T-stage or M-state). We focused on tumor progression rather than on comparing cancerous with healthy samples. This strategy allowed us to concentrate on 7 clinical transitions (a-g) instead of seeking lncRNA profiles that signify cancer initiation. Importantly, while hundreds of Tr-lncRNAs were detected across all transitions, a substantial fraction was unique to specific cancer types and mostly to a single one. Focusing on lncRNAs that exhibit pan-cancer roles exposes lncRNAs that are strongly coordinated across numerous cancer types, as shown for XIST and USP9Y (Fig. 5).

When Tr-lncRNA expression trends were compared, the uniqueness of each cancer type was revealed. Specifically, non-overlapping pattern between cancers, suggesting a context-specific regulatory process. This view further supports the role of lncRNAS in determining chromatin state, lineage-specific transcription factors and tumor microenvironment cues (Ransohoff et al., 2018). This high cancer type specificity stands in contrast to the generality of protein-coding oncogenes or tumor suppressors (e.g., KRAS, TP53, PTEN). According to such high specificity, the Tr-lncRNAs that signify the transition to metastasis (M0 → M1) may serve as biomarkers with prognostic and therapeutic relevance. The signature of Tr-lncRNAs in specific cancers that switch to a metastatic state motivates their use for follow-up in clinical and experimental contexts (Statello et al., 2021). The cancer-specific nature of lncRNA programs supports the development of stage-specific diagnostics. Additionally, when we combine all identified Tr-lncRNAs from this study (any lncRNA that appeared in any of the cancer-transition analyses (total 2,399), 13.3% of Tr-lncRNAs predicted to localize in exosomes (lncBook (Ma et al., 2019)), therefore liquid biopsy might be used as a promising avenue for lncRNA-based non-invasive cancer monitoring.

The fraction of antisense (AS) among the lncRNA collection in TCGA is substantial. This finding is in accord with the dependency of lncRNAs’ expression by their genomic positions (Engreitz et al., 2016). In our effort to assign functional relevance, we observed that among TR-lncRNAs, the fraction of AS reach 47% (based on the relaxed definition from lncBook). This enrichment is consistent with the contribution as cis-regulatory elements acting in transcriptional interference and as promoter-enhancer scaffold (Fig. 8). In studying cancer induction genomic aberrations and mutations in driver genes dictate cancer (Hanahan, 2022). In contrast, the expression of lncRNAs explores other aspects of cancer regulation such as checkpoints, chromatin regulation, nuclear scaffold all are related to cancer transition stages. We propose a platform that highlights known and previously overlooked lncRNAs as candidates for experimental validation in cancer-relevant model systems.

Nonetheless, this study also uncovers several limitations that warrant future investigation. Technically, dealing with lncRNAs is quite sensitive to the protocol used for data normalization (see Methods). To avoid the biases stemming from low expression levels that define most lncRNAs, we focus our analysis primarily on z values >3, which indicate overexpression during cancer progression. For the Tr-lncRNAs with negative values (statistical threshold of z-values <-3), the number of Tr-lncRNA candidates is lower (e.g., Fig. 4C), likely resulted from the bounded by the minimal expression level requested for sample consistency. Additionally, most lncRNAs have multiple transcripts (Fig. 9C), many of which are alternative spliced. In this study, we reduced the complexity of lncRNA transcripts by considered only a single representative. Lastly, the low overlap between cancer types, while emphasizing specificity, raises challenges for the generalizability of our findings for clinical use. As the data from TCGA reports on bulk RNA-seq, the ability to dissect cell-type contributions to the lncRNA signals is minimal. Single-cell resolution combined with spatial transcriptomics becomes attractive directions to test whether Tr-lncRNAs arise from the tumor cells themselves or reflect the TME with a contribution of infiltrated immune cell populations (Armand et al., 2021). Recently, machine learning approach was developed toward immune-related lncRNA signature in cancer (Li et al., 2023). We expect that these approached will benefit from a benchmark of cancer specific refined set of Tr-lncRNAs. By illuminating the dynamic and context-specific behavior of lncRNAs across disease stages and cancer types, our study lays the groundwork for lncRNA-guided diagnostics and therapeutics.

## Methods

### lncRNA collection and nomenclature

We have used the comprehensive, non-redundant, gene-centric database of human non-coding RNAs (ncRNAs) from GeneCaRNA (Barshir et al., 2021). This collection was constructed through the integration of gene centric resources (HGNC, NCBI Gene, and Ensembl), in addition to the rich transcript-centric view from RNAcentral database. Altogether, the GeneCaRNA reported on ∼220, 000 unique ncRNA identifiers. While, GeneCaRNA database refers to the >22k genes that overlapped with gene-centric resources, additional ∼188k were novel entries based on Transcript-inferred GeneCaRNA Genes (TRIGGs) (Barshir et al., 2021).

Naming of lncRNAs is often associated with specific catalog and database. The major resources of lncRNAs use non coherent indexing systems. Additional resources such as Lnc2Cancer 3.0 (Gao et al., 2021), LncRNADisease (Bao et al., 2019), TANRIC (Li et al., 2015) and LNCipedia (Volders et al., 2019) were included in GeneCaRNA protocol mapping. For consistency, we have adapted the genes names from GeneCaRNA according to genomic position of the lncRNA (i.e., considering the transcription directionality). To secure unique mapping, we revisited cases of ambiguity and required that >90% sequence overlap >200 nt as a minimal mapping requirement. We have ignored classification by RNA structure (i.e., linear, circular) (Ahadi, 2021) or by subcellular localization.

While often the lncRNA names are used as technical identifiers, some prefix and suffix are informative. For example, Linc01234 refer to Long intergenic noncoding RNA number 1234), TTTY refers to Testis-specific transcript, Y-linked. FAM refers to Family with sequence similarity, AF are indicative of being provisional or temporary. In addition, a suffix of AS and HG denote the potential position of an antisense lncRNA and long host gene for miRNAs, respectively. An early effort to create a comprehensive catalog and unified naming protocol of lncRNAs reported for ∼57.5k genes from GENCODE V.18 (Yan et al., 2015).

### Expression and clinical data for major cancer samples

RNA sequencing (RNA-seq) data and the associated clinical annotations were obtained from The Cancer Genome Atlas (TCGA) via the Genomic Data Commons (GDC) data portal (Tomczak et al., 2015a). The RNA-seq data were generated using the HiSeq platform, employing a poly(A)+ selection protocol and paired-end sequencing strategy. Raw read counts and normalized expression values (FPKM) for annotated lncRNAs were extracted and processed. In TCGA, the reported measures are provided as RSEM (RNA-seq by expectation xaximization) for estimating gene and isoform expression levels from the raw data. The normalization is based on log2[FPKM+1] to avoid biases from extremely small numbers.

Clinical metadata including cancer type, patient age, sex, tumor stage, and survival outcomes were curated from the Pan-Cancer Atlas of TCGA, ensuring consistent stage classification across studies. We used the routine clinical classification as captured by the TNM categories. We consider the T classes (tumor stage) and M, the presence or absence of metastasis. We do not address the involvement of regional lymph nodes (N) due to lack of robust classification. For the T-stages, we consider stages I to IV as the stages mark greater local invasion (and included T2a and T2b as stage II, when available). Stage grouping followed the AJCC (American Joint Committee on Cancer) criteria supports the merging of Tumor (T) stage I and II. In this study we have not included samples marked as T0 (normal, no evidence of a primary tumor) or Tis (carcinoma in situ). The M0 is defined by not having distant metastasis while M1 indicates distant metastasis present.

### Dimensionality reduction

For the normalized expression data, we performed t-SNE analysis (t-distributed stochastic neighbor embedding) to visualize the data. Each point represents a sample of the analyzed tumor. The spatial proximity reflects the overall similarity in the expression profiles of the listed lncRNAs. Using sklearn t-SNE package to project samples on 2D space for visualization using data frame, with samples (rows) and the expression of each of the relevant lncRNAs (columns).

### Differentially expressed transitional lncRNAs

Two types of labeling were used for the clinical annotations. The T and M system. For the T-stage, we applied each sample to its stage (I-IV) and for the M (metastasis) we determine the M0, as primary tumor and M1 as metastasis. Each lncRNA that was expressed in any of the major cancer types (see list Supplemental **Table S2**) was calculated to meet a z-value >|3|. The calculation of the z-value is based on considering the mean and standard deviation of the differences of any specific cancer.

In order to capture the significance, magnitude and directionality of a change in lncRNA expression in specific transition in a cancer type (i.e I➔II in BRCA), we first filtered the lncRNAs to include only lncRNAs who has an expected count value of 1.0 in at least 30% percent of the samples (in at least one of the two stages of the transition) in order to avoid small fractional expression values likely to reflects noise. Then, we used z-scores over the log2(FC) of the means over the log(x+1) expression values of the lncRNA across samples in the two stages participating in the transition. This was calculated from the mean value of expression (in any specific cancer type) over the samples of a selected stage relative to the consecutive one (e.g., the difference of stage T-II to stage T-I). Substracting the expression in a form of log_2_(FC) value from the precalculated expected counts of the analyzed cancer type (marked as log(x+1)). We confirmed that the log2(FC) approximate a normal distribution.

To further test the robustness of the identified Tr-lncRNAs, we inverted the log values back to the expected counts (log(x+1)->x) and taking the mean over these values and only then apply the log(mean+1) and calculating z-scores over the log2fold changes. By clipping extreme samples (5%) we observed strong consistency with the mean over log values method as described above.

### Annotation enrichment

We analyzed the annotations of the lncBook 2.0 resource (Li et al., 2023). The annotations for conservation and subcellular location are according to lncBook 2.0 definitions. We also adhered to lncBook genomic definitions and assigned lncRNAs according to strand directionality. For example, the Intronic (AS) lncRNAs are transcribed from the AS strand of protein-coding genes, with their sequences are within an intron. The overlapping (AS) lncRNAs contain coding genes within an intron on the AS strand. The AS lncRNAs are transcribed from the AS strand of protein-coding genes, and the entire sequences of lncRNAs are covered by protein-coding genes, or intersect partially with them. The protein capacity annotation relied on data from

Ribo-seq and mass spectrometry (MS) evidence. Notably, there are often multiple small protein associated with a lncRNA gene with protein capacity. Statistical measures for any feature (total 143) are determined by the Mann-Whitney test and Chi-square for numerical and binary features, respectively. A threshold of FDR <0.05 was set for statistically significant enrichment.

## Data Availability

Data processing protocols, code, visualizations and large tables are available in the GitHub. The Supplemental Tables S1-S4 and Figure S1-S6v are available by the Journal online information. https://github.com/Daniel-Behar/Transitional-lncRNA-Signatures-Reveal-Distinct-Stages-of-Cancer-Progression-and-Metastasis

### Declaration of competing interest

The authors declare no competing interests. The authors report no conflict of interest.

### Funding

This study was supported in part by the the Center for Interdisciplinary Data Science Research (CIDR, 3035000440).

## Acknowledgments

We thank the member of the Linial’s lab for their support throughout this work, with a special thanks to Roei Zucker and Keren Zohar. We thank the GeneCard team (Weizmann Institute of Science) for providing the GeneCaRNA mapping protocol that combined and harmonized many lncRNA resources.

## Notes

https://github.com/Daniel-Behar/Transitional-lncRNA-Signatures-Reveal-Distinct-Stages-of-Cancer-Progression-and-Metastasis

### Competing Interest Statement

The authors have declared no competing interest.

