## Supplemental Figure S1-s6 for "Transitional lncRNA Signatures Reveal Distinct Stages of Cancer Progression and Metastasis"

<u>**Supplemental Materials**</u>

**Supplemental Fig. S1.** 2-D visualization of t-SNE for healthy tissues

**Supplemental Fig. S2.** 2-D visualization of t-SNE for 4 T-stages (I to IV)

**Supplemental Fig. S3.** Distribution of samples over T-stages and M-state over cancer types

**Supplemental Fig. S4.** Distribution of expression by the z score for example of Stage I→II for BRCA.

***Supplemental Fig. S5.*** *Similarity matrix of Tr-lncRNAs for major cancer types for 6 diffeent transitions.*

***Supplemental Fig. S6.*** *A network connectivity for ME to ML a transition*

**Supplemental Table S1.** *major cancer types abbreviation and statistics*

**Supplemental Table S2.** *Pan cancer analysis (PC) for Tr-lncRNA with ≥2 cancer types (E→L and ME→ML)*

**Supplemental Table S3.** Matrix of all lncRNA, for all cancer types, for z-threshold for a-g transitions

**Supplemental Table S4.** Feature enrichment for 143 lncRNA features

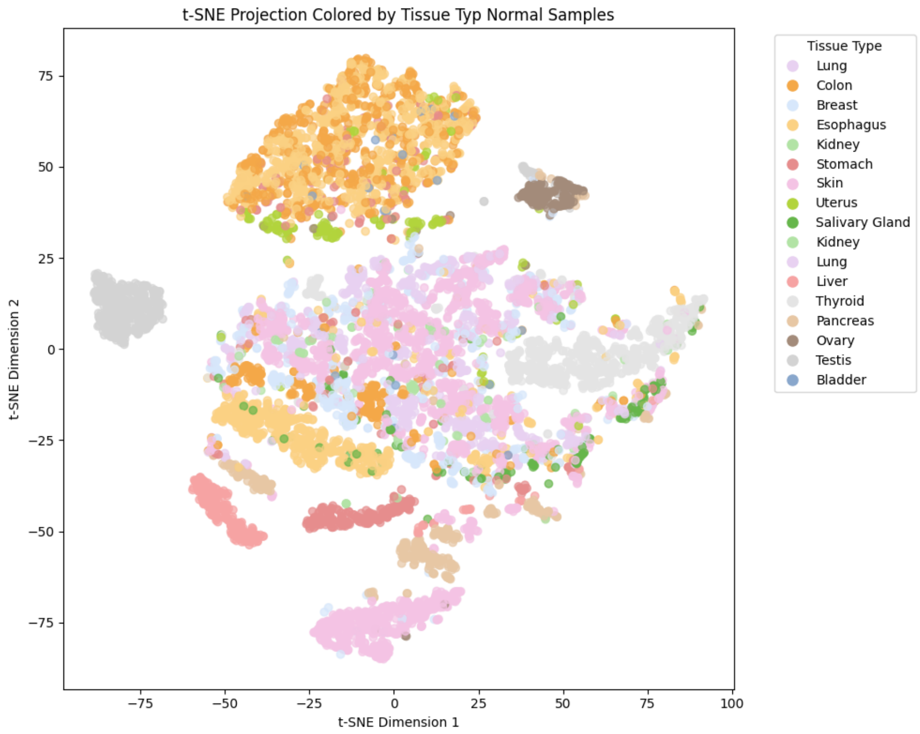


**Figure S1.** Clustering by t-SNE from all listed lncRNA by the RNA-seq. Data was extracted from GTEx tissues.

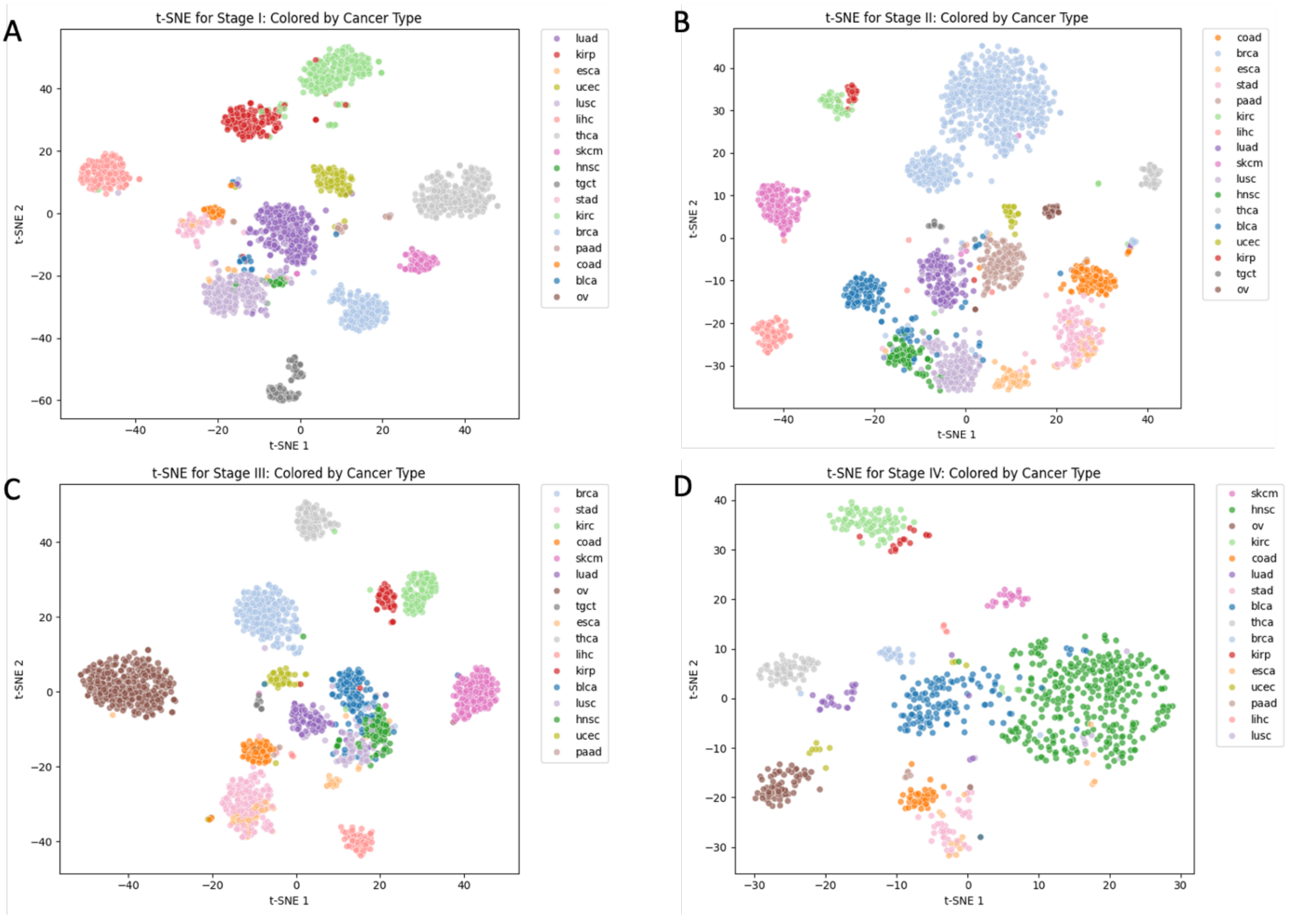

**Figure S2.** Cancer type clustering by t-SNE by lncRNAs expression profiles. The color coded for the 2D-embedding is shown for the cancer types (total 16) and for the T-stages. **(A)** Stage I, **(B)** Stage II, **(C)** Stage III**,** and **(D)**, Stage IV**.** The expression data used included all lncRNAs that were expressed and normalized in the analyzed samples. The resulted clusters are colored by cancer types as indicated in TCGA for each sample. Note that sample sizes differ substantially between stages. For example, while there were only a few OV samples in stages I and II, the majority of the OV samples are labeled at stage III.

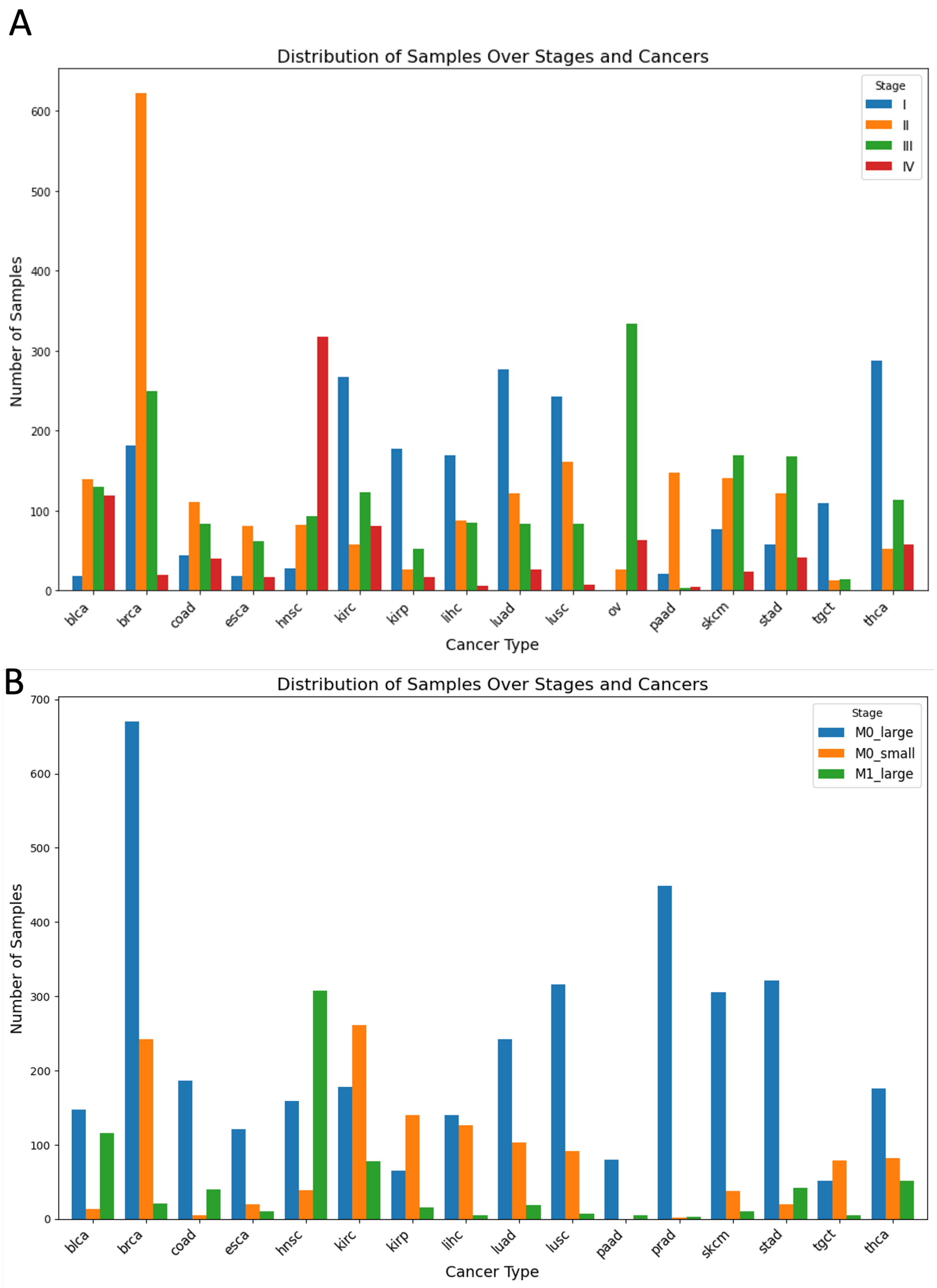


**Figure S3.** Statistics of Tr-lncRNAs by cancer types. **(A)** Sample view partitioned among the 16 major cancer types according to their T-stage (I-IV), sorted alphabetically. **(B)** Sample view partitioned among the 16 major cancer types according to their M-state (M0/M1), sorted alphabetically. Note that OV only labelled by T-stage and PRAD only by M-state.

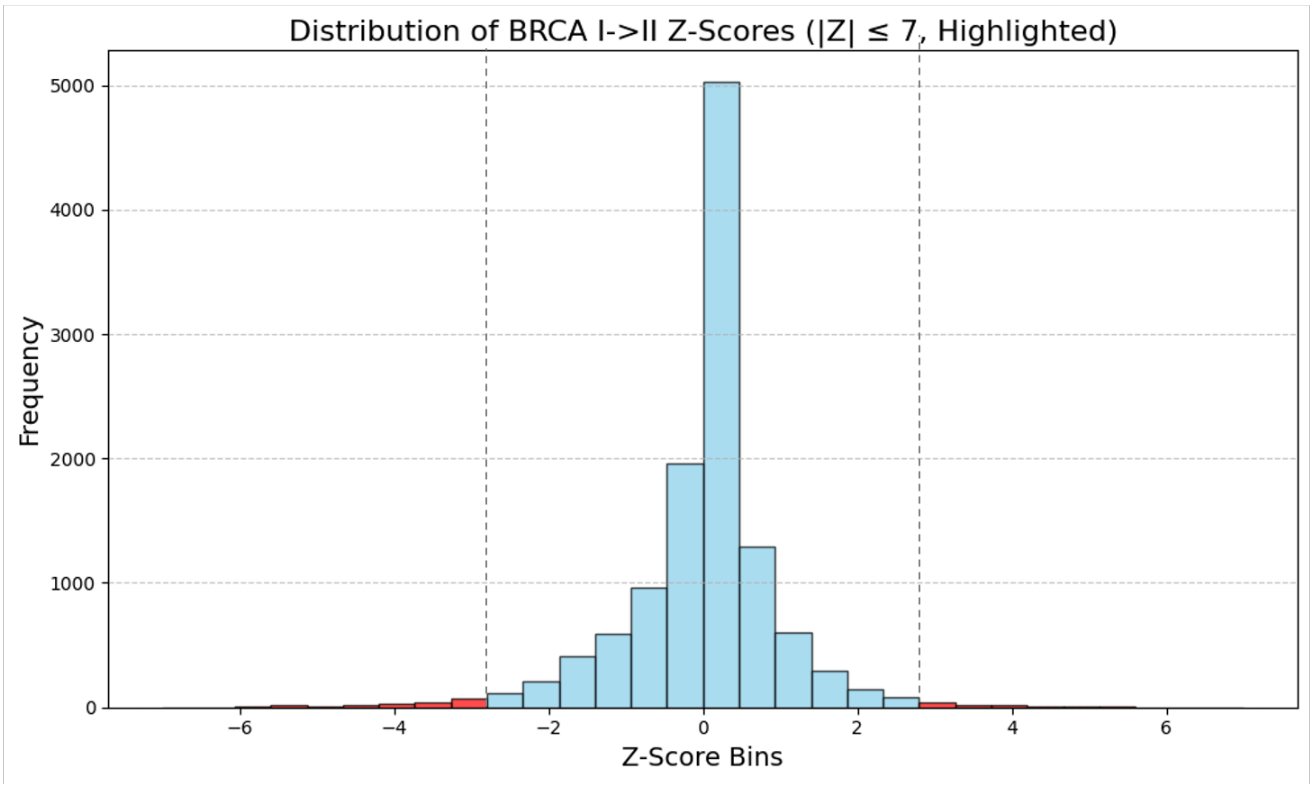


**Figure S4.** Distribution by the z-score is illustrated for BRCA transition I → II. The threshold of z-value >|3| was applied for the rest of the analysis. only tails of the distribution were further analyzed. There are a few examples with z-value >|7| that were trimmed for the visualization clarity.

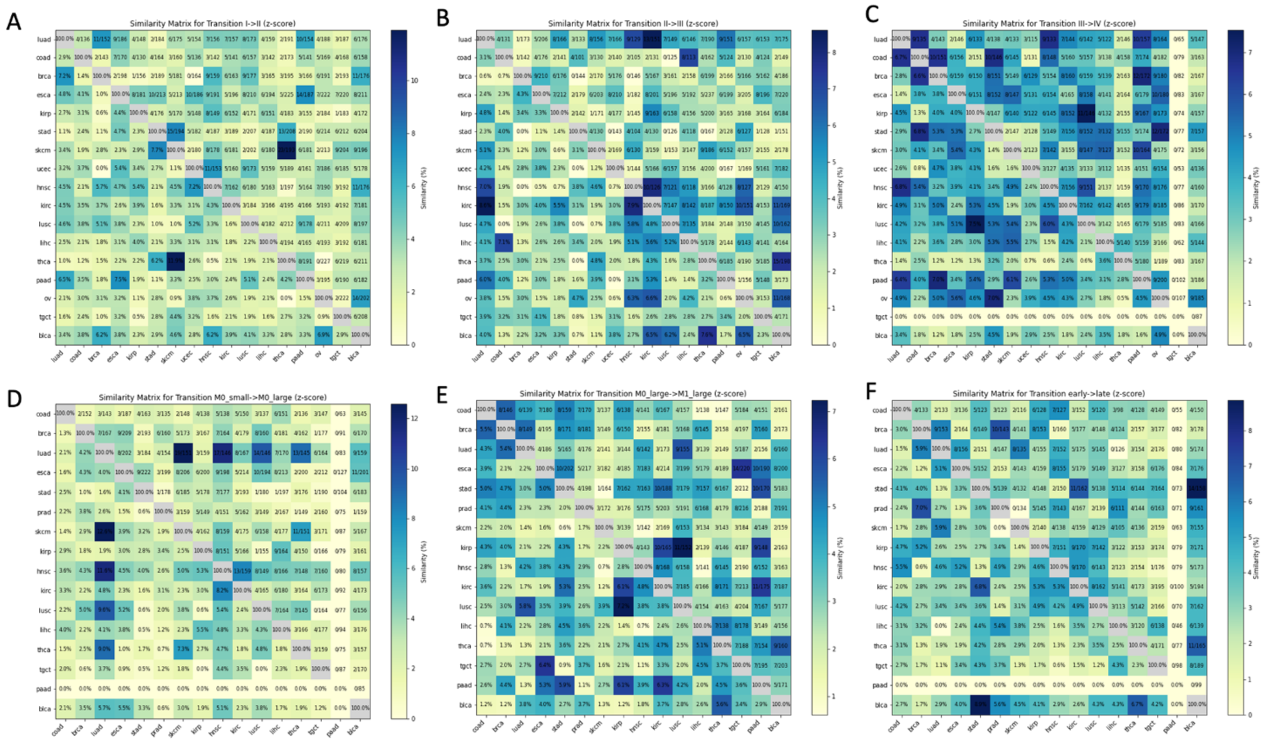

**Figure S5.** Heatmaps of the similarity matrices of Tr-lncRNAs among all major cancer types from TCGA. The color (yellow to dark blue) marks the percentage of sharing with any other cancer types (bottom left), and in a symmetrical position above the diagonal, the actual number of shared Tr-lncRNAs from the total number of potential lncRNAs. **(A)** Heatmap for the T-stage I→II transition (transition a) on 17 cancer types. **(B)** Heatmap for the T-stage II→ III transition (transition b) on 17 cancer types. **(C)** Heatmap for the T-stage III→ IV transition (transition c) on 17 cancer types (note that TCGT failed to provide reliable results). **(D)** Heatmap for the M-state M0_small→M0_large transition e) on 16 cancer types. Note that PAAD failed to provide information for this transition. **(E)** Heatmap for the M-state M0_large→M1_large transition f) on 16 cancer types. **(F)** Heatmap for the M-state ME→ML transition g, on 16 cancer types. Note that PAAD failed to provide information for this transition.

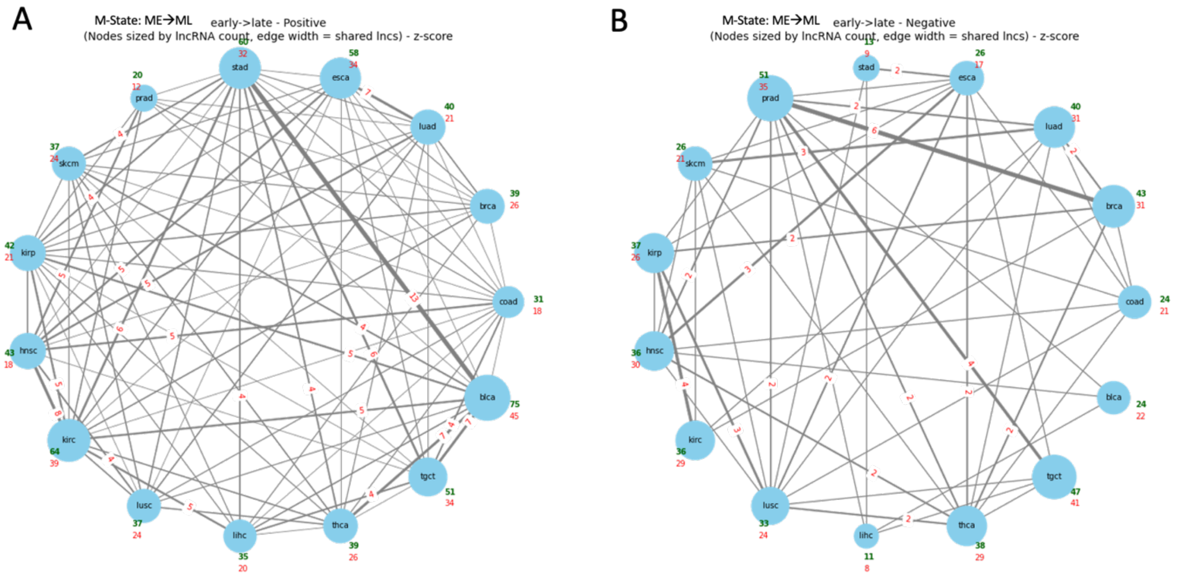


**Figure S6.** Connectively map of all cancer types for M-state. **(A)** The nodes mark the cancer types. The transition of M-state ME→ML (transition g) for the positive z-values. **(B)** The nodes mark the cancer types. The transition of M-state ME→ML (transition g) for the negative z-values. The number of shared lncRNAs are indicated on the edges The size of the node reflects the number of TR-lncRNA in each cancer type. The width of the edges indicates the number of shared Tr-lncRNAs (marked by numbers on the edge). In red fonts are the number of unique Tr-lncRNAs for each cancer type.
